# Single-cell Total-RNA Profiling Unveils Regulatory Hubs of Transcription Factors

**DOI:** 10.1101/2023.12.14.571592

**Authors:** Yichi Niu, Jiayi Luo, Chenghang Zong

**Author notes:** Corresponding Author: Chenghang Zong. These authors contributed equally to this work (Yichi Niu, Jiayi Luo).

## Abstract

Recent development of RNA velocity uses master equations to establish the kinetics of the life cycle of RNAs from nascent RNA to mature RNA to degradation. To feed this kinetic analysis, simultaneous measurement of nascent RNA and mature RNA in single cells is greatly desired. However, the majority of single-cell RNA-seq chemistry only captures mature RNA species to measure gene expressions. Here, we develop a one-step total-RNA chemistry-based scRNA-seq method: snapTotal-seq. We benchmarked this method with multiple single-cell RNA-seq assays in their performance in kinetic analysis of cell cycle by RNA velocity. Next, with LASSO regression between transcription factors, we identified the critical regulatory hubs mediating the cell cycle dynamics. We also applied snapTotal-seq to profile the oncogene-induced senescence and identified the key regulatory hubs governing the entry of senescence. Furthermore, from the comparative analysis of nascent RNA and mature RNA, we identified a significant portion of genes whose expression changes occurred in mature RNA but not to the same degree in nascent RNA, indicating these gene expression changes are mainly controlled by post-transcriptional regulation. Overall, we demonstrate that snapTotal-seq can provide enriched information about gene regulation, especially during the transition between cell states.

---

The rapid development of single-cell RNA-seq (scRNA-seq) has enabled large-scale characterization of different cell states, which provides valuable information about the changes that occurred during the cell-state transitions, such as development and differentiation processes. Following this characterization, it is desired to decode the underlying regulatory mechanisms. Here, we reason that this regulatory information could be derived based on the comparative analysis between nascent RNA and mature RNA. However, the majority of scRNA-seq methods, including the high-throughput platform of 10x chromium, are mature RNA only (oligo-dT) based methods^1–5^, making them less ideal for the comparative analysis due to inefficient detection of nascent RNA species^6^. Meanwhile, metabolic labeling strategies have been incorporated into the mature-RNA based scRNA-seq pipelines to label the newly synthesized RNA^7–11^. But, this approach demands live cells, and the tracing ability is limited to the time following the addition of nucleoside analogs. In addition, the potential side effects on cell physiology caused by nucleoside analogs should also be taken into consideration. Here, we showed that the total-RNA based scRNA-seq methods without the need for chemical labeling are more suited for kinetic and regulatory analyses.

So far, three main single-cell total-RNA methods have been developed in recent years, including MATQ-seq^12^ by Sheng *et al.* in 2017, Smart-seq-total method^13^ by Isakova *et al.* in 2021, and VASA-seq by Salmen *et al.* in 2022^14^. Total-RNA-seq chemistry has also been applied to profile the transcriptome of individual neuronal synaptosomes^15^. Despite the advantage of the single-cell total-RNA sequencing approach for RNA velocity analysis^16,17^, the chemistries of current total-RNA-based methods are generally more complicated than mature RNA based methods, especially in comparison to SMART-seq chemistry^4,18,19^. Here, by combining multiple annealing chemistry allowed by MALBAC primers^20^ and the template switching chemistry^4,18,19,21^ (**Supplementary** Fig. 1), we developed a one-step single-cell total-RNA-seq chemistry. We refer to this new assay as snapTotal-seq, which can be easily implemented on liquid-handling platforms.

Next, we benchmarked the performance of snapTotal-seq, SMART-seq3, CEL-Seq2, SMART-seq-total, and VASA-seq in their ability to capture the cell cycle dynamics through RNA velocity analysis. As a result, in comparison to mature RNA based methods, total-RNA based methods demonstrated substantial improvement in recapitulating the transcriptional dynamics of the cell cycle, with snapTotal-seq achieving the best performance.

With the trajectory data from RNA velocity, we showed that by LASSO regression of the nascent RNA expressions of a transcription factor (TF) against the mature RNA expressions of the rest of TFs, we identified the important TF hubs that mediate the cell state transitions during the cell cycle. Furthermore, the comparative analysis between nascent RNA and mature RNA expression also revealed the substantial role of post-transcriptional regulation in the expression of a significant portion of cell cycle genes.

Besides the cell cycle, we further expanded the application of snapTotal-seq to investigate the regulatory network in oncogene-induced senescence (OIS). As a result, we also successfully identified the key regulatory components that orchestrated cell state transition during OIS. Overall, these findings underscore the ability of snapTotal-seq to characterize the transcription/splicing/degradation dynamics of RNAs and, with greater significance, to identify the key transcription hubs driving cell-state transitions.

## Results

### Chemistry of snapTotal-seq

The overall chemistry of snapTotal-seq is shown in **Fig. 1a**. After the lysis of single cells; we used MALBAC primers^20^ to initiate the reverse transcription at low temperature, which allows the random but efficient primer binding to multiple sites on RNAs, therefore warranting the total-RNA detection ability. Next, we gradually ramped up the temperature to promote the reverse transcription. Once the reverse transcription stopped at the sites with difficult secondary structures or reached the 5’ end of RNA, template-switching then occurred with the MALBAC primers serving as the template-switching oligos. In total, we performed ten cycles of multiple annealing and extension (without melting steps), during which successful template switching generated cDNA amplicons. With the special design of MALBAC primers, the primer dimer is not detectable in the final product even when the annealing steps reach as low as 8°C. After cDNA amplification, cell-specific barcodes were introduced to the amplified products by a double-strand conversion step. Next, the cells with different barcodes were pooled together for the library construction (**Supplementary** Fig. 2). It is worth noting that all the reads in our method have UMI sequences.

**Figure 1.**
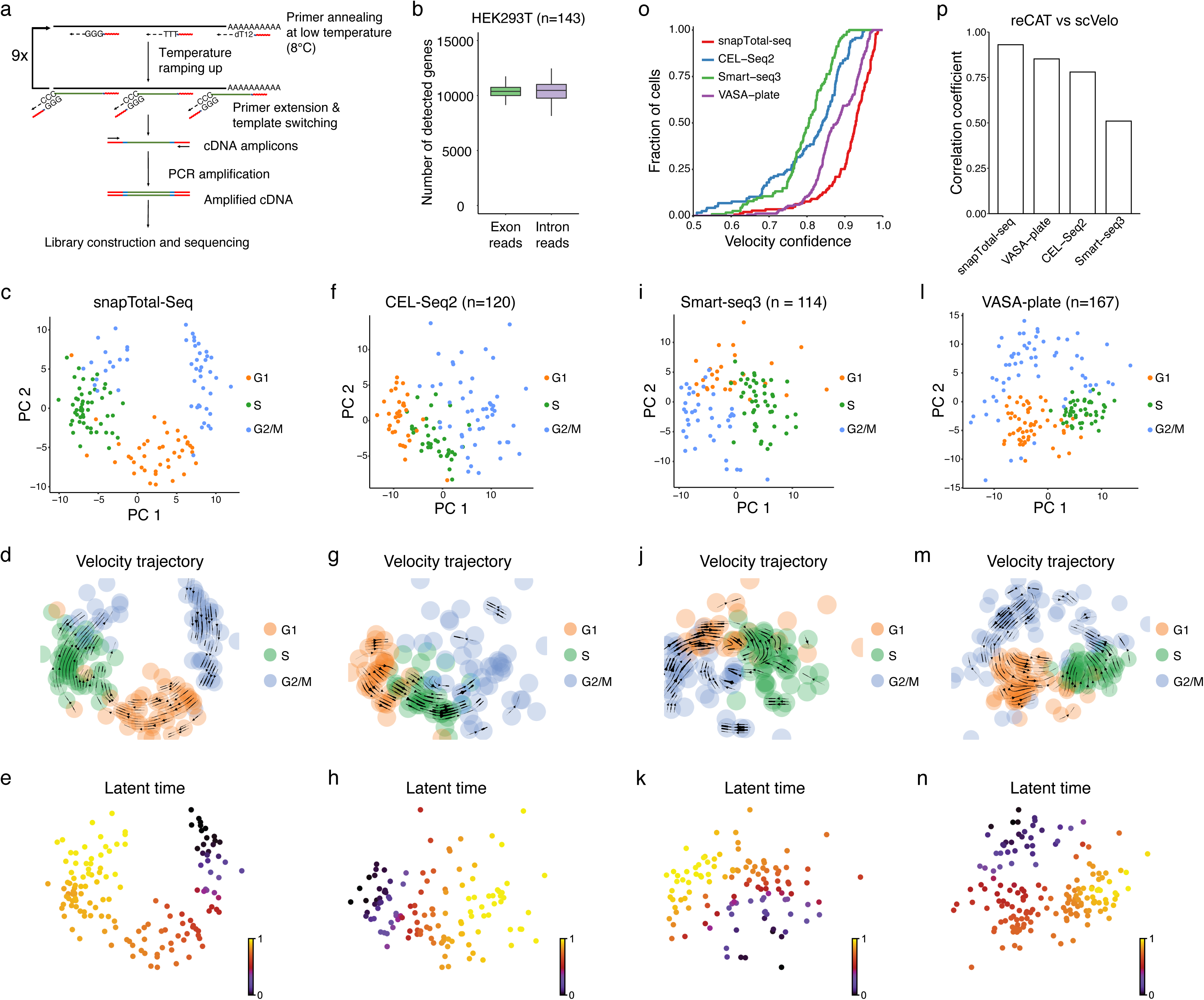
Development of snapTotal-seq and benchmark analysis. (**a**) Schematic of snapTotal-seq. (**b**) The number of detected genes by exon reads, or intron reads in single HEK293T cells by snapTotal-seq. (**c**) PCA plot of the HEK293T cells. The cells are colored by their corresponding cell cycle phases. (**d**) The velocity trajectory projected by RNA velocity analysis. (**e**) Latent time inferred by RNA velocity. (**f**) PCA plot of the HEK293T cells sequenced by CEL-Seq2. (**g**) The projected velocity trajectory of the HEK293T cells sequenced by CEL-Seq2. (**h**) Latent time inferred by RNA velocity for CEL-Seq2. (**i**) PCA plot of the HEK293T cells sequenced by Smart-seq3. (**j**) The projected velocity trajectory of the HEK293T cells sequenced by Smart-seq3. (**k**) Latent time inferred by RNA velocity for Smart-seq3. (**l**) PCA plot of the HEK293T cells sequenced by VASA-plate. (**m**) The projected velocity trajectory of the HEK293T cells sequenced by VASA-plate. (**n**) Latent time inferred by RNA velocity for VASA-plate. (**o**) The velocity confidence scores of the HEK293T cells sequenced by different methods. (**p**) The correlation coefficients between the reCAT based cell cycle trajectory and the RNA velocity based cell cycle trajectory achieved by different methods.

We first performed the cross-species experiment using HEK293T cells and 3T3 mouse cells. We observed that the crosstalk between the two species was rare (**Supplementary** Fig. 3a). Next, we performed two batches of HEK293T cells. We showed that the technical batch effects were minimal as the single cell data of different batches were completely overlapped in the principal component analysis (PCA) (**Supplementary** Fig. 3b). Overall, we were able to detect 10372±518 genes based on exon reads and 10431±875 genes based on intronic reads with ∼1 million uniquely mapped reads per cell (**Fig. 1b**), which confirms the effective capture of both mature RNA and nascent RNA by snapTotal-seq.

### Benchmarking comparison of single-cell RNA-seq methods in RNA velocity analysis

The primary advantage of total-RNA based methods lies in their ability to provide the data that facilitate the modeling of gene expression processes, from transcription to splicing and RNA decay, as described by the master equations of the RNA velocity^16,17^. We first applied RNA velocity analysis to the snapTotal-seq data. As shown in **Fig. 1c-e**, we observed that the velocities clearly captured the circular dynamic transitions between different cell cycle phases (G2/M -> G1 -> S -> G2/M) with high confidence scores (0.93±0.073, **Supplementary** Fig. 3c). To verify the result of RNA velocity analysis, we applied another algorithm: reCAT^22^ to analyze the cell cycle dynamics using the same data. Different from *ab initio* based analysis by RNA velocity, reCAT determines the cell cycle dynamics based on the known cell cycle genes. The pseudotime derived from reCAT is shown in **Supplementary Fig.3d**. And the latent time from RNA velocity and the pseudotime from reCAT were highly correlated with Pearson’s coefficient larger than 0.9 (**Supplementary Fig.3e**).

Next, we performed a benchmark comparison between snapTotal-seq and four scRNA-seq methods: Smart-seq3^4^, CEL-Seq2^5^, VASA-seq^14^ (plate version) and Smart-seq-total^13^. Among them, Smart-seq3 and CEL-Seq2 employ oligo-dT-based reverse transcription strategy, while VASA-seq and Smart-seq-total are recently developed total-RNA based scRNA-seq methods. Firstly, we observed that the gene detection rate of snapTotal-seq is similar to VASA-seq based on either exon reads or intronic reads. And they are substantially higher than the two dT-based methods (**Supplementary** Fig. 4a-b). Interestingly, we noticed Smart-seq-total has a low gene detection with either exon or intron reads, which is potentially due to the dominance of detection of miscRNA in their data^13^, suggesting that this method is more suited for detecting short RNA species. Due to this significant detection difference, we excluded Smart-seq-total from the downstream RNA velocity analyses.

Next, we performed PCA analysis and RNA velocity analysis on each dataset. We observed that the connections between different cell cycle states were not well positioned for both dT-based methods in the PCA plots (**Fig. 1f-k**). In contrast, both snapTotal-seq (**Fig. 1c-e**) and VASA-seq (**Fig. 1l-n**) showed a clear circular distribution describing different cell cycle phases. As a result, the velocity confidence scores of Smart-seq3 and CEL-Seq2 were significantly lower than those of snapTotal-seq and VASA-seq. We also noticed that snapTotal-seq has a better confidence score than VASA-seq (**Fig. 1o**). Meanwhile, we also employed reCAT analysis to infer the cell cycle trajectory based on the expression of known cell cycle genes (**Supplementary** Fig. 4c-e). As a result, our method achieved the highest correlation coefficient between the results of the unsupervised and supervised analysis (**Fig. 1p**). Based on the systematic comparison described above, we concluded that snapTotal-seq outperformed other methods in characterizing the gene expression kinetics.

### Comparative analysis between nascent and mature RNAs identified three types of cell cycle genes (CCGs)

Based on the constructed cell cycle trajectory as described above, we then performed the trajectory-based differential gene expression analyses for both mature RNA and nascent RNA using tradeSeq^23^. We identified 1325 genes with significant changes at the mature RNA level (FDR < 0.1) and 962 genes with significant changes at the nascent RNA level (FDR < 0.1) along the cell cycle (**Fig. 2a** **& Table S1**).

**Figure 2.**
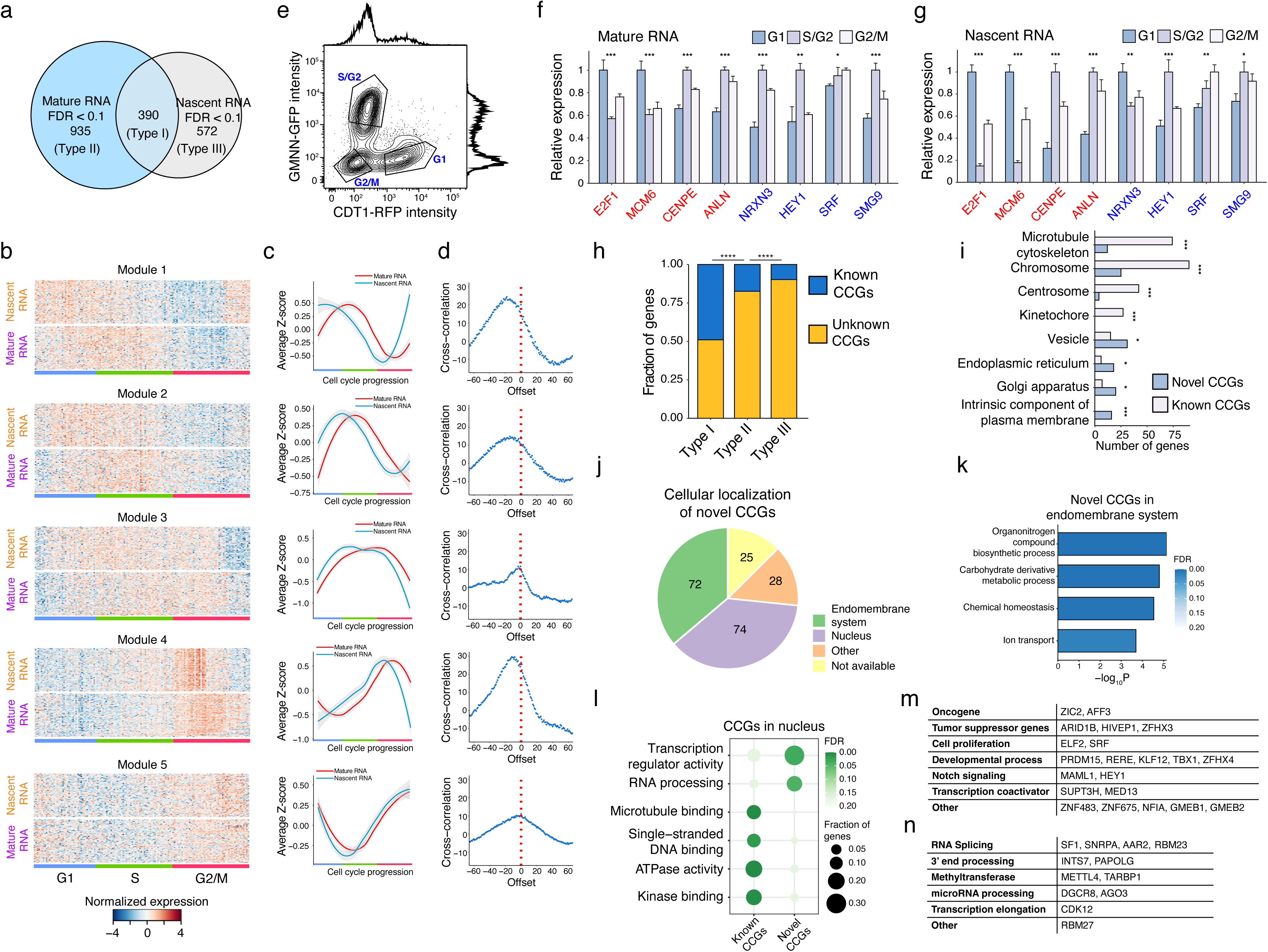
Functional analysis on Type I CCGs. (**a**) Differentially expressed genes identified by tradeSeq at the mature RNA or the nascent RNA level along the cell cycle in HEK293T cell line. Three types of CCGs are defined. Type I CCGs are the genes with significant expression changes in both nascent RNA and mature RNA along the cell cycle. Type II CCGs are the genes with significant expression changes in only mature RNA along the cell cycle. Type III CCGs are the genes with significant expression changes in only nascent RNA along the cell cycle. (**b**) The gene expression heatmap of Type I CCGs at the mature RNA and nascent RNA levels along the cell cycle. The genes were clustered into 5 kinetic modules based on their transcriptional dynamics along the cell cycle. (**c**) The smoothed gene expression curves along the cell cycle trajectory for 5 kinetic modules. The smoothed curves were derived by using Loess function. Shade, 0.95 confidence interval. Z-score is used to define the gene expression changes. (**d**) The cross-correlation between the expression curves of the nascent RNA and mature RNA. The cross-correlation was calculated by using numpy.correlate function in Python. (**e**) The flow cytometry analysis on the HEK293T cells with FUCCI markers. (**f-g**) qRT-PCR validation of the differential expression of 8 CCGs at the mature RNA level (**f**) or at the nascent RNA level (**g**) in different cell cycle phases. The known CCGs are labeled in red, and the novel CCGs are labeled in blue. Student’s t-test, * p < 0.05, ** p < 0.01, *** p < 0.001, **** p < 0.0001. (**h**) The proportions of novel CCGs in three types of CCGs. The list of known CCGs is compiled by including the gene list of the cell cycle pathway in the Gene Ontology database, the gene list of the cell cycle pathway in the Reactome database, and the gene list from Cyclebase. Fisher’s exact test, * p < 0.05, ** p < 0.01, *** p < 0.001, **** p < 0.0001. (**i**) Differential enrichment in different cellular compartments between the known CCGs and novel CCGs. Fisher’s exact test, * p < 0.05, ** p < 0.01, *** p < 0.001, **** p < 0.0001. (**j**) The cellular localization of all novel CCGs. (**k**) The Gene Ontology (GO) enrichment of the novel CCGs localized in the endomembrane system. (**l**) The GO enrichment of the known CCGs and the novel CCGs localized in the nucleus. (**m**) The functional classification of the transcription factors belonging to novel CCGs. (**n**) The functional classification of the RNA processing factors belonging to novel CCGs.

We noticed that there are 390 genes that showed significant changes in both nascent and mature RNA, which we denoted as Type I CCGs, and 935 genes that showed only significant changes in mature RNA, which we denoted as Type II CCGs, and 572 genes that showed only significant changes in nascent RNA, which we denoted as Type III CCGs. For Type II CCGs, the lack of significant changes in nascent RNA suggests that gene expression changes that occurred in the mature RNAs are mainly contributed by post-transcriptional regulation (**Supplementary** Fig. 5a-b). In comparison, gene expression changes in Type I CCGs are mainly contributed by transcriptional regulation. For a large number of genes in the Type III CCGs, it is interesting to see the transcriptional variations that occurred to these genes are effectively buffered out at the mature RNA level, likely also through post-transcriptional regulation-based mechanisms, which is worth future investigation.

### Identification of five kinetic modules and novel genes in Type I CCGs

For Type I CCGs, we identified five kinetic modules (**Fig. 2b** **& Supplementary** Fig. 5c-g). As shown in **Fig. 2b-c**, a clear time delay occurs between the changes in nascent RNA and mature RNA, which mainly corresponds to the splicing process. We quantified the coupling between the dynamics of nascent RNA and mature RNA by the cross correlation between two expression curves (**Fig. 2d**). The close coupling confirms that the changes in the expression of these genes mainly originated from the regulation at the transcription step.

To validate the detected differentially expressed genes (DEGs) by snapTotal-seq along the cell cycle, we used HEK293T cells with the expression of FUCCI markers^24^ to collect the cells at G1, S/G2 and G2/M phases (**Fig. 2e**). We then randomly chose eight genes from the detected Type I CCGs and performed qRT-PCR to measure their gene expression levels. As a result, the differential expression of these genes at different cell cycle stages was confirmed for both nascent RNA and mature RNA (**Fig. 2f-g**).

Besides the genes with known functions in cell cycle regulation (i.e., known CCGs), we observed that over 50% of Type I CCGs (199 out of 390) had not been associated with cell cycle regulation previously (here we refer to them as novel CCGs) (**Fig. 2h** **& Supplementary** Fig. 5h). In contrast to known CCGs, which are significantly enriched with mitosis related structures, the novel CCGs are significantly enriched in the endomembrane system, including endoplasmic reticulum, Golgi apparatus, and intrinsic component of plasma membrane (**Fig. 2i-j**). Functional enrichment shows that these genes are involved in the synthesis of organonitrogen compounds, metabolism of carbohydrate derivatives, ion transportation, and maintenance of chemical homeostasis (**Fig. 2k**).

Next, the second major subset of novel CCGs (74 out of 199) reside in the nucleus (**Fig. 2j**). In contrast to the known CCGs in the nucleus that are enriched with microtubule binding, single-strand DNA binding and ATPase activity, the novel CCGs are enriched in transcriptional regulators and RNA processing factors (**Fig. 2l**). The related pathways of these novel CCGs are shown in **Fig. 2m** for transcriptional regulators and **Fig. 2n** for RNA processing factors. The identification of the abundant novel cell cycle genes directly shows the advantage of our method in transcriptional kinetics analysis.

### Identification of TF hubs regulating the five kinetic modules along the cell cycle

Next, we sought to identify the transcription factors (TFs) that could govern the five kinetic modules observed in Type I CCGs. To do so, we utilized LASSO regression^7^ to identify the associations between the mature RNA changes of TFs and the nascent RNA changes of the rest of the genes in different modules (**Fig. 3a**). It is worth noting that, in comparison to oligo-dT-based methods, the simultaneous detection of both nascent RNA and mature RNA by our method allows us to directly link the expression of TFs (mature RNA levels) to the transcriptional dynamics of their target genes (nascent RNA levels) through this analysis. As a result, we identified 23 TFs with their potential downstream genes being significantly enriched in Type I kinetic modules (**Fig. 3b**). Furthermore, based on the transcriptional activities of these 23 TFs (i.e., the changes in the nascent RNA levels of their target genes), we successfully identified 4 TF hubs (**Fig. 3c**). The correlation matrix between these hubs is shown in **Fig. 3d**.

**Figure 3.**
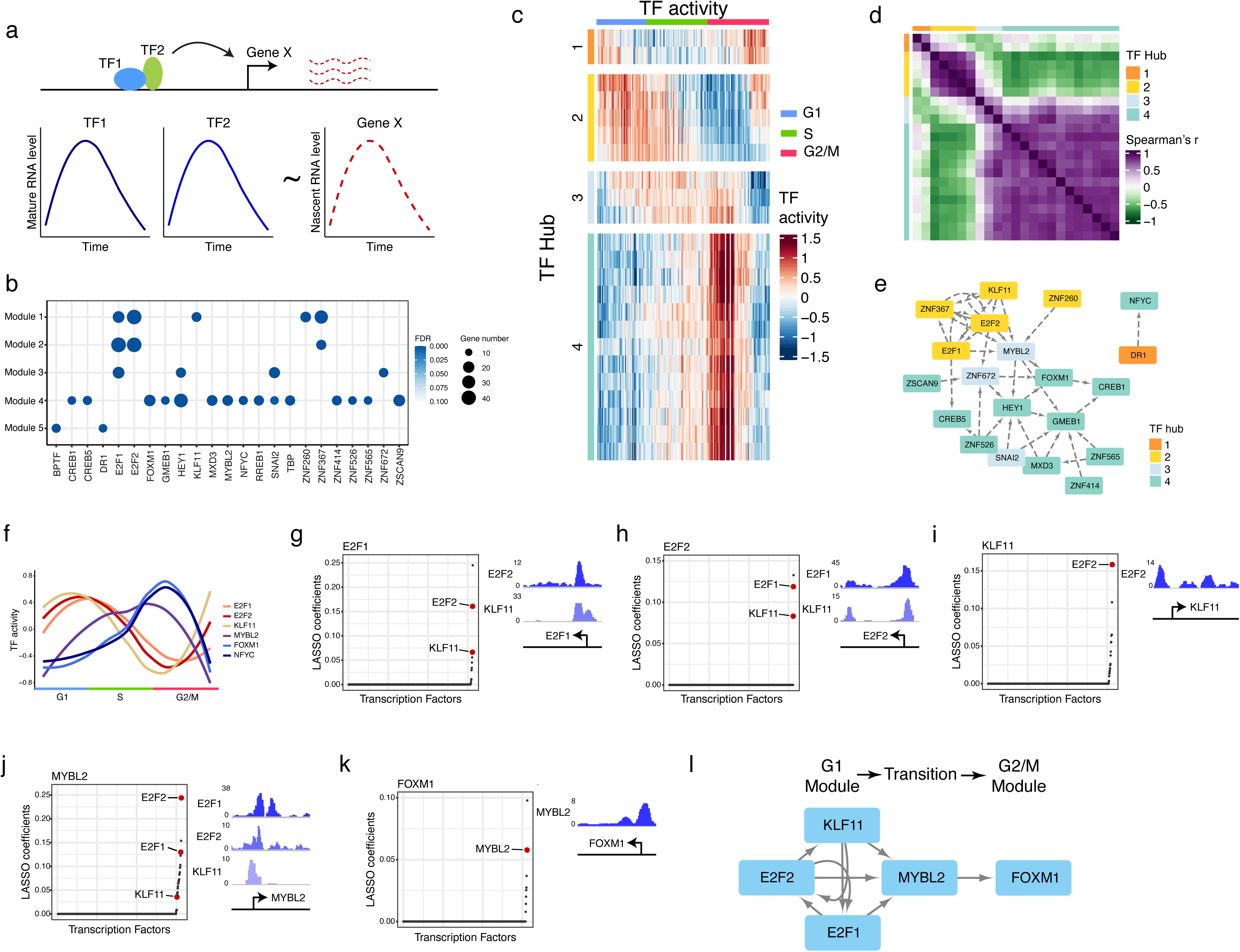
Reconstructing TF regulatory network of the cell cycle. (**a**) Schematic outlining the approach used to establish the regulatory links between TFs and their target genes. (**b**) TFs whose associated genes are significantly enriched in Type I kinetic modules. (**c**) Heatmap of the activities of the TFs with significant enrichment with Type I kinetic modules. (**d**) Heatmap of the correlation coefficients between the activities of each pair of TFs. (**e**) TF association network established based on the TF-TF links identified with LASSO regression. (**f**) The activities of the verified TFs along the cell cycle. The smoothed curves were derived by using Loess function. (**g-k**) Establish the regulatory links between different TFs by correlating the changes in the nascent RNA of the TF of interest with the changes in the mature RNA of the other TFs. The direct regulatory links (colored in red) were identified by ChIP-seq verification. The corresponding ChIP-seq peaks were plotted on the right. (**l**) TF regulatory network established based on the direct regulatory links between different TFs.

Next, to identify the potential connections between these 23 TFs, we performed the LASSO regression between the changes in the nascent RNA of a TF and the changes in the mature RNA of the rest of the expressed TFs using LASSO regression. This analysis produces the TF association network as shown in **Fig. 3e**. Interestingly, we noticed that Hub 3 (light blue gene blocks), while only composed of 3 genes, was located in the middle of Hub 2 (yellow gene blocks) and Hub 4 (cyan gene blocks), and was connected to the genes in both Hub 2 and Hub 4, indicating that Hub 3 play important roles in mediating the transition from G1/S to G2/M state.

### Validation of TF regulations based on ChIP-seq data and Motif analysis

From the LASSO regression inferred TF hubs, we next evaluate the direct regulation based on the published ChIP-seq datasets^25^ or motif analysis^26^ (**Methods**). Among the candidate TFs, the ChIP-seq data was unavailable for 12 of them. Out of the 11 TFs with ChIP-seq data, we identified 6 TFs (*E2F1*, *E2F2*, *KLF11, FOXM1, NFYC* and *MYBL2* ) whose downstream genes showed significant enrichment with the binding targets identified by ChIP-seq (FDR < 0.05, **Supplementary** Fig. 5i) or displayed a significant enrichment of the corresponding motif sequence at their promoter regions (normalized enrichment score (NES) > 3, **Table S2**), confirming the direct regulatory relationships between these TFs and their downstream genes identified by LASSO regression.

Among the 6 TFs verified by ChIP-seq data, *E2F1*, *E2F2* and *KLF11* TFs have been known for their roles in regulating G1/S transition^27,28^, and consistently, the transcriptional activities of the associated gene modules were clearly increased in G1 and early S phases (**Fig. 3f**). The regulation relation between *KLF11, E2F1* and *E2F2* is shown in **Fig. 3g-i**. For both *FOXM1* and *NFYC*, their associated gene modules were highly activated in late S and early G2 phases (**Fig. 3f**). Consistent with this observation, *FOXM1* is the well-known master regulator of G2/M phase^27,29,30^. Next, we observed that the *MYBL2*, another major regulator of G2/M gene expression^31,32^, was clearly activated prior to the activation of *FOXM1* module (**Fig. 3f**), implying that *MYBL2* functions as an intermediate TF that could play important roles in the transition from G1/S to G2/M state. It is worth noting that *MYBL2* is located in Hub 3. The detailed regulation relation between *MYBL2* and *KLF11, E2F1*, *E2F2* and *FOXM1* are shown in **Fig. 3j-k**.

Overall, we observed that the G1/S TFs formed a positive feedback loop to promote G1/S phase gene expression (**Fig. 3l**). Consistent with previous findings by Skotheim *et al*.^33^, the positive feedback loops ensure the stability of G1/S state. Interestingly, our data unveiled that all of these G1/S TFs positively regulated the expression of *MYBL2* (**Fig. 3j** **& 3l**). The upregulated *MYBL2* subsequently activated the expression of *FOXM1* (**Fig. 3k** **& 3l**), leading to the activation of G2/M phase gene expression. This observation shows the pivotal role of *MYBL2* in driving the cell state transition from G1/S towards G2/M. To further verify the generality of this observation, we sequenced another two cell lines (U2OS and HPNE) using snapTotal-seq, and similar observations were obtained (**Supplementary** Fig. 6-7).

In summary, our results demonstrated the ability of snapTotal-seq to recapitulate the TF regulatory network that drives the cell-state transition. For the other 5 TFs that lack significant enrichment in the ChIP-seq data, it could be caused by either the binding targets revealed by ChIP-seq being cell-line specific or the identified downstream genes being indirect targets of these TFs. Future studies are required to further investigate the regulatory roles of these TFs in cell cycle progression.

### Identification of novel CCGs under post-transcriptional regulation

Type II CCGs were featured by their significant changes at the mature RNA level but not at the nascent RNA level (**Supplementary** Fig. 5a-b), indicating the involvement of post-transcriptional regulation of these genes. Based on their dynamic changes along the cell cycle, we also identified five kinetic modules (**Fig. 4a-b**). The cross-correlation analysis shows that the coupling between the gene expression changes in nascent RNA and mature RNA is not as significant as Type I CCGs (**Fig. 2d** and **Fig. 4c**). Gene ontology (GO) enrichment analysis was then carried out on each kinetic module. Module 2 has significant enrichment with DNA replication and repair pathways. Modules 1, 4, and 5 were significantly enriched with pathways that are not directly associated with cell cycle progression (**Fig. 4d**). These new pathways include RNA metabolism, RNA processing, vesicle transportation, etc. These results suggest that the activities of many biological processes are coordinated with cell cycle progression by post-transcriptional regulations.

**Figure 4.**
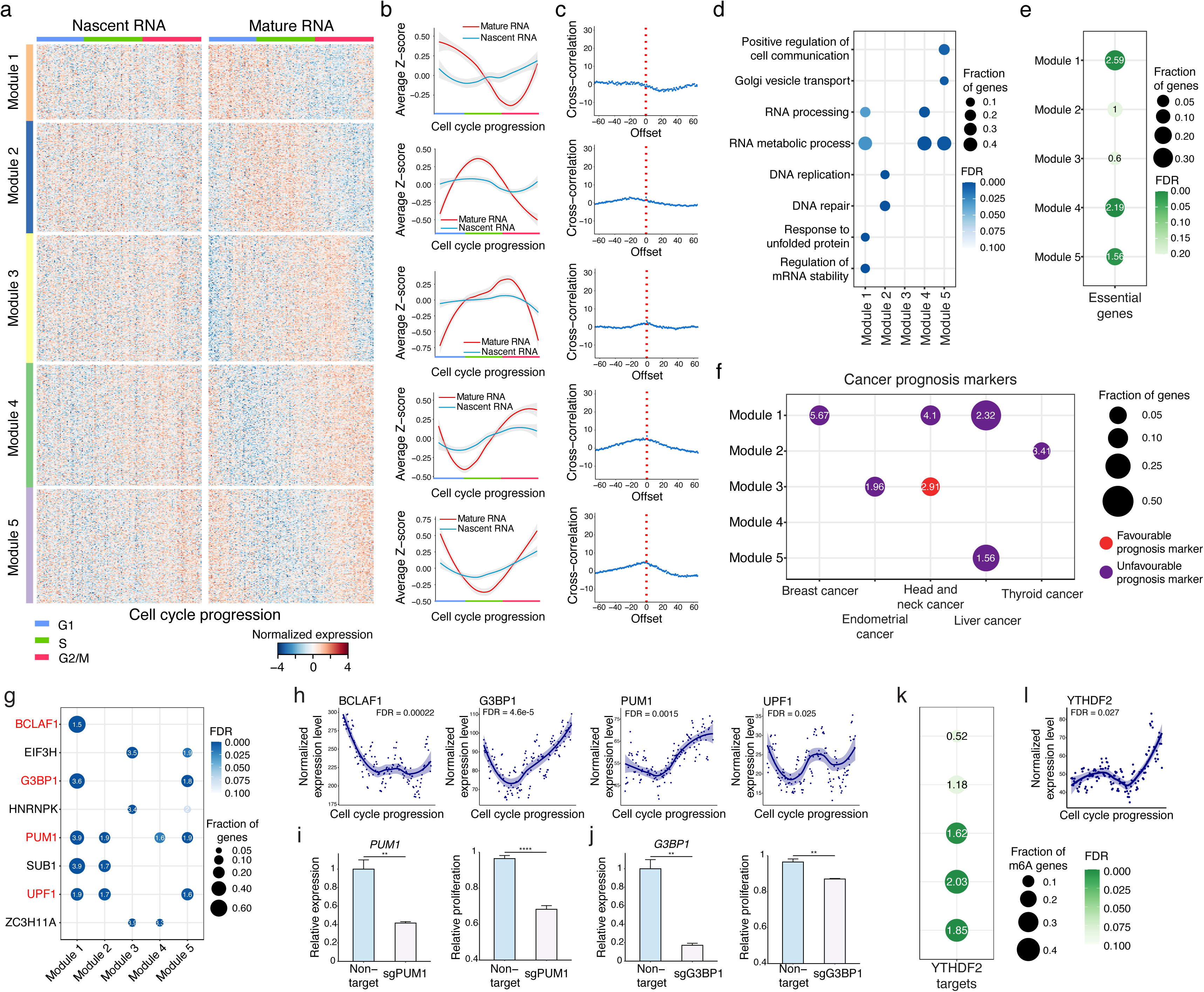
Post-transcriptional regulation in Type II CCGs. (**a**) The gene expression heatmap of Type II CCGs at the mature RNA and nascent RNA levels along the cell cycle. The genes were clustered into 5 kinetic modules based on the expression patterns of their mature RNA along the cell cycle. The periodic expression of these genes is clearly observed at the mature RNA level, while the expression patterns at the nascent RNA level are much less distinct. (**b**) The smoothed gene expression curves along the cell cycle trajectory for 5 kinetic modules. The smoothed curves were derived by using Loess function. Shade, 0.95 confidence interval. Z-score is used to define the gene expression changes. (**c**) The cross-correlation between the expression curves of the nascent RNA and mature RNA. The cross-correlation was calculated by using numpy.correlate function in Python. We did not observe a clear coupling between the expression changes in nascent RNA and mature RNA. (**d**) The GO enrichment in 5 kinetic modules of Type II CCGs. (**e**) The significant enrichment of pan-essential genes in different gene modules. The significance of enrichment is determined by Fisher’s exact test, and the enrichment fold in each module is labeled. The known CCGs in each module are excluded from this analysis. (**f**) The significant enrichment (FDR < 0.1) of cancer prognostic markers in different gene modules. The significance of enrichment is determined by Fisher’s exact test, and the enrichment fold in each module is labeled. The known CCGs in each module are excluded from this analysis. (**g**) The significant enrichment (FDR < 0.1) of the target genes of the RNA binding proteins in Type II CCGs. The significance of enrichment is determined by Fisher’s exact test, and the enrichment fold in each module is labeled. (**h**) The gene expression dynamics of the four RNA binding proteins along the cell cycle. The smoothed curves were derived by using Loess function. Shade, 0.95 confidence interval. (**i**) The cell proliferation was significantly decreased in *PUM1* knockdown cells compared to the cells infected with the non-target sgRNA. (**j**) The cell proliferation was significantly decreased in *G3BP1* knockdown cells compared to the cells infected with the non-target sgRNA. Student’s t-test, * p < 0.05, ** p < 0.01, *** p < 0.001, **** p < 0.0001. (**k**) The significant enrichment of the target genes of YTHDF2 in the Type II CCGs with m6A modification. The significance of enrichment is determined by Fisher’s exact test, and the enrichment fold in each module is labeled. (**l**) The gene expression dynamics of *YTHDF2* along the cell cycle. The smoothed curves were derived by using Loess function. Shade, 0.95 confidence interval.

To evaluate the important roles of the novel Type II CCGs in the cell cycle, we examined the essentiality of these genes by using the genetic screening datasets from Project Achilles^34^. As a result, we found that the pan-essential genes were significantly overrepresented among the novel CCGs in Modules 1, 4, and 5 (**Fig. 4e**). This observation indicates that the functions of these post-transcriptionally regulated kinetic modules are important for cell proliferation. Next, we explored whether these genes play critical roles in cancer progression since cancer is a disease of dysregulation of the cell cycle. Interestingly, we found that except for Module 4, the novel CCGs in the rest of the modules were significantly enriched with the prognostic markers of different cancer types^35^, further indicating that cancer cells likely need to alter post-transcriptional regulations to fit with the abnormal cell proliferation (**Fig. 4f**).

### Multiple post-transcriptional regulation mechanisms underlying Type II CCGs

Next, we investigated the potential mechanisms that drive the differential expression of Type II CCGs. Considering that RNA binding proteins (RBPs) have been reported as a class of proteins regulating the fate of RNA at different post-transcriptional processing steps^36–38^, we examined their potential roles in regulating the differential expression of Type II CCGs. Among the known 150 RBPs whose binding targets have been thoroughly examined^39^, 22 RBPs were differentially expressed along the cell cycle in our data.

To evaluate their functions in cell cycle progression, we then compared their binding targets to Type II CCGs. As a result, we identified 8 RBPs whose binding targets were significantly enriched in at least one Type II CCG module (**Fig. 4g**). It is worth pointing out that 4 out of these 8 RBPs (*UPF1*, *BCLAF1*, *PUM1,* and *G3BP1*) have already been reported to regulate RNA stability and decay^39^. We indeed observed that the expression patterns of these 4 RBPs along the cell cycle were consistent with the expression patterns of their target gene groups (**Fig. 4h**).

Next, to further validate our findings, we knocked down two non-essential genes: *PUM1* and *G3BP1* (**Fig. 4i-j, Supplementary** Fig. 8a). We did not test the knockouts of *UPF1* and *BCLAF1* since they have been classified as pan-essential genes^34^. As a result, we observed that the cell growth was significantly decreased in *PUM1* knockdown cells (**Fig. 4i**), which suggests that *PUM1* indeed plays a critical role in regulating cell proliferation in the HEK293T cell line. The knockdown of *G3BP1* also led to decreased cell proliferation with statistical significance (**Fig. 4j**).

It is also worth noting that we identified 14 genes whose binding targets were not significantly enriched in Type II CCGs. Interestingly, these genes are mainly composed of splicing factors, rRNA processing factors, and miRNA processing factors (**Supplementary** Fig. 8b). For the splicing factors, the lack of target enrichment in Type II CCGs suggests that they regulate the general splicing process to adapt to different cell cycle phases or modulate periodic alternative splicing along cell cycle, which is consistent with previous studies ^40,41^.

Besides RBPs, N6-methyladenosine (m6A) modification, one of the most abundant modifications on mammalian mRNA, has also been shown to regulate the fate of RNA by recruiting different readers^42–44^. One of the well-known readers of m6A modification is YTHDF2, and it has been shown to affect the RNA stability^45,46^. Here, we observed that YTHDF2 target genes^46^ are significantly enriched in the genes with m6A modification^47,48^ in Module 3, 4, and 5 (**Fig. 4k**), and meanwhile, the gene expression of *YTHDF2* is also cell-cycle dependent (**Fig. 4l**). These results showed that RNA modification contributes to the regulation of Type II CCGs. Overall, our analysis suggested that RNA binding proteins and m6A modification of RNA play important roles in regulating the expression of Type II CCGs along the cell cycle.

### RNA velocity analysis of oncogene-induced senescence

To further test the ability to dissect the gene expression kinetics and gene regulation underlying cell state transition, we applied snapTotal-seq to characterize the gene regulation in the entry into the oncogene-induced senescence (OIS). Here, we utilized the HPNE cells with inducible KRAS^G12D^ expression^49^. After the activation of KRAS^G12D^, we collected the cells on Days 1, 2, 3, and 5 (**Fig. 5a**), and in total, we sequenced 642 cells using snapTotal-seq. The cells were plotted on the UMAP and were labeled by the time points (**Fig. 5b**) and the assigned cell cycle stages (**Fig. 5c**). Interestingly, we observed that the cells are distributed in two clusters. One cluster (top right cluster in **Fig. 5c**) is mainly composed of G1, S, and G2/M phases, while the other one (bottom left cluster in **Fig. 5c**) is mainly composed of the G0 population based on reCAT analysis. By comparing the percentage of different cell cycle stages at each time point, we observed a gradual increase of the G0 cells from day 1 to day 5 (**Supplementary** Fig. 9a), suggesting that the cells continuously entered the G0 phase following the induction of oncogenic KRAS.

**Figure 5.**
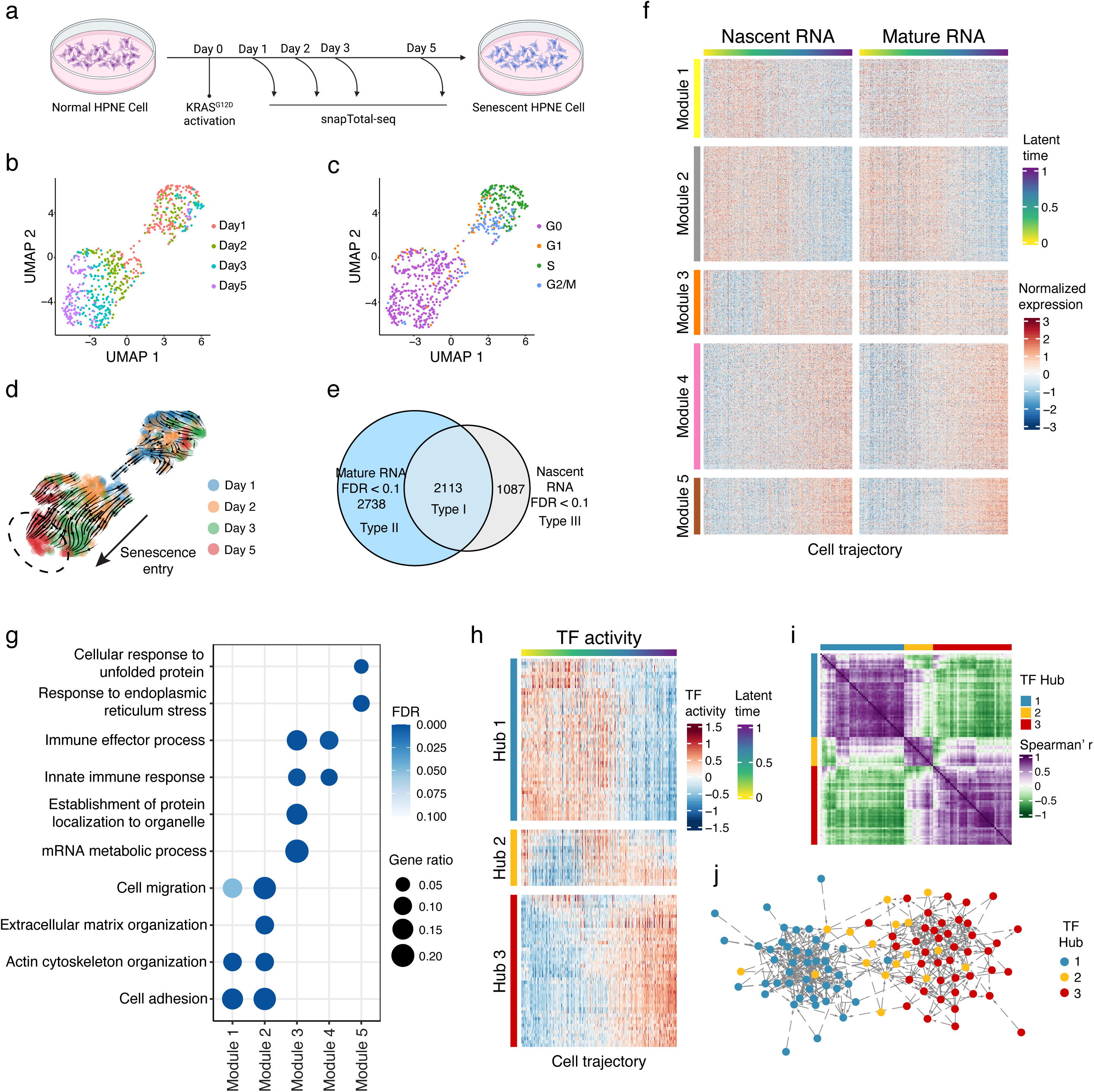
Transcriptional dynamics during oncogene-induced senescence. (**a**) Experimental scheme for studying gene expression dynamics of oncogene-induced senescence. (**b-c**) UMAP visualization of the HPNE cells with KRAS^G12D^ overexpression. The cells are colored by the time points (**b**) or the cell cycle stages (**c**). (**d**) The projected velocity trajectory of oncogene-induced senescence by RNA velocity analysis. The cells are colored by the time points. (**e**) Differential gene expression analyses identified the genes with significant changes at both mature RNA and nascent RNA levels (Type I), only at the mature RNA level (Type II), and only at the nascent RNA level (Type III) along oncogene-induced senescence. (**f**) The gene expression heatmap of the genes with significant changes at both mature RNA and nascent RNA levels (Type I DEGs) along the latent time. (**g**) The GO enrichment in 5 kinetic modules of Type I DEGs. (**h**) Heatmap of the activities of all enriched TFs along cell trajectory. (**i**) Heatmap of the correlation coefficients between the activities of each pair of TFs. (**j**) TF association network established based on the TF-TF links identified with LASSO regression.

Next, we applied RNA velocity analysis to construct the trajectory of OIS. We observed a clear trajectory within the G0 population indicated by the bold arrow in **Fig. 5d**. The trajectory follows the time order of day 1, day 2, day 3, and day 5 (**Fig. 5d** and **Supplementary** Fig. 9b). Therefore, the bottom-left region of the G0 cluster (dashed circle in **Fig. 5d**) corresponds to the endpoint: the senescent state. The successful trajectory inference was also confirmed by the velocity confidence scores (0.88±0.050) (**Supplementary** Fig. 9c).

### Comparative analysis between nascent and mature RNAs along OIS

Next, to investigate the gene expression dynamics during OIS, we performed the differential gene expression analysis for both mature RNA and nascent RNA based on the trajectory within the G0 population established by RNA velocity analysis (**Fig. 5e** **& Table S3**). Similar to cell cycle analysis, we define the genes with significant changes in both mature RNA and nascent RNA as Type I DEGs, the genes with only significant changes in mature RNA as Type II DEGs, and the genes with only significant changes in nascent RNA as Type III DEGs. In total, we identified 2113 genes as Type I DEGs, 2738 genes as Type II DEGs, and 1087 genes as Type III DEGs (**Fig. 5e**).

### Identification of five gene expression kinetic modules in OIS

We focused on the transcriptional regulation of the Type I DEGs during OIS. Through an analysis of their transcriptional dynamics, we first detected five kinetic modules among Type I DEGs (**Fig. 5f-g**). We noticed that Module 1 and 2 were significantly downregulated along the senescence entry, and they were significantly enriched with the genes involved in cell adhesion and extracellular matrix organization (**Fig. 5f-g** **& Supplementary** Fig. 9d). Conversely, the expression level of Module 3 was transiently downregulated at the early stage, followed by a rapid rebound to its original level (**Fig. 5f** **& Supplementary** Fig. 9e). Lastly, the expression of Modules 4 and 5 demonstrated significant upregulation along the trajectory, underscoring their pivotal roles in establishing OIS (**Fig. 5f** **& Supplementary** Fig. 9f). Specifically, Module 4 that is enriched with the genes involved in the immune response (**Supplementary** Fig. 9f), was quickly activated after the induction of oncogenic stress, followed by a linear increase in their expression levels. In contrast, the proteotoxic stress response pathway (i.e., Module 5, **Supplementary** Fig. 9f) exhibited an initial slow activation, which was succeeded by a rapid elevation in expression levels at the middle time point.

### Identification of TF Hubs underlying the kinetic modules by LASSO regression

To identify the TF hubs that regulate these kinetic modules, we established the association between TFs and their target genes based on the LASSO analysis, the same as analyzed in the cell cycle (**Fig. 3a**). In total, we identified 112 TFs whose linked genes were significantly enriched with Type I kinetic modules (FDR < 0.01). These TFs were then grouped into 3 distinct regulatory hubs based on their activities along the trajectory of OIS (**Fig. 5h-i**). We observed that Hub 1 is mainly associated with kinetic modules 1 and 2, Hub 2 is mainly associated with kinetic modules 3, and Hub 3 is mainly associated with kinetic modules 4-5.

Next, we identify TF-TF associations within the TF hubs or between the TF hubs using LASSO analysis. In **Fig. 5j**, we colored TFs based on their hubs as determined above. As a result, we observed that TFs from the same hub tend to form densely interconnected sub-networks (Student’s t-test, p < 2.2e-16 for all TF hubs). More interestingly, we observed that Hub 2 is located between Hub 1 and Hub 3 (**Fig. 5j**), indicating their important roles in the shifting process from one regulatory controller to another regulatory controller.

### Validation of specific regulations based on ChIP-seq data and Motif analysis

To further pin down the regulatory relationships between the TFs, we examined whether associated genes were significantly enriched with either the ChIP-seq validated binding targets or the gene promoters carrying the corresponding binding motif. As a result, we found 25 TFs with significant enrichment (**Table S4**). We then labeled these 25 verified TFs based on their respective TF hubs described above (**Fig. 6a**).

**Figure 6.**
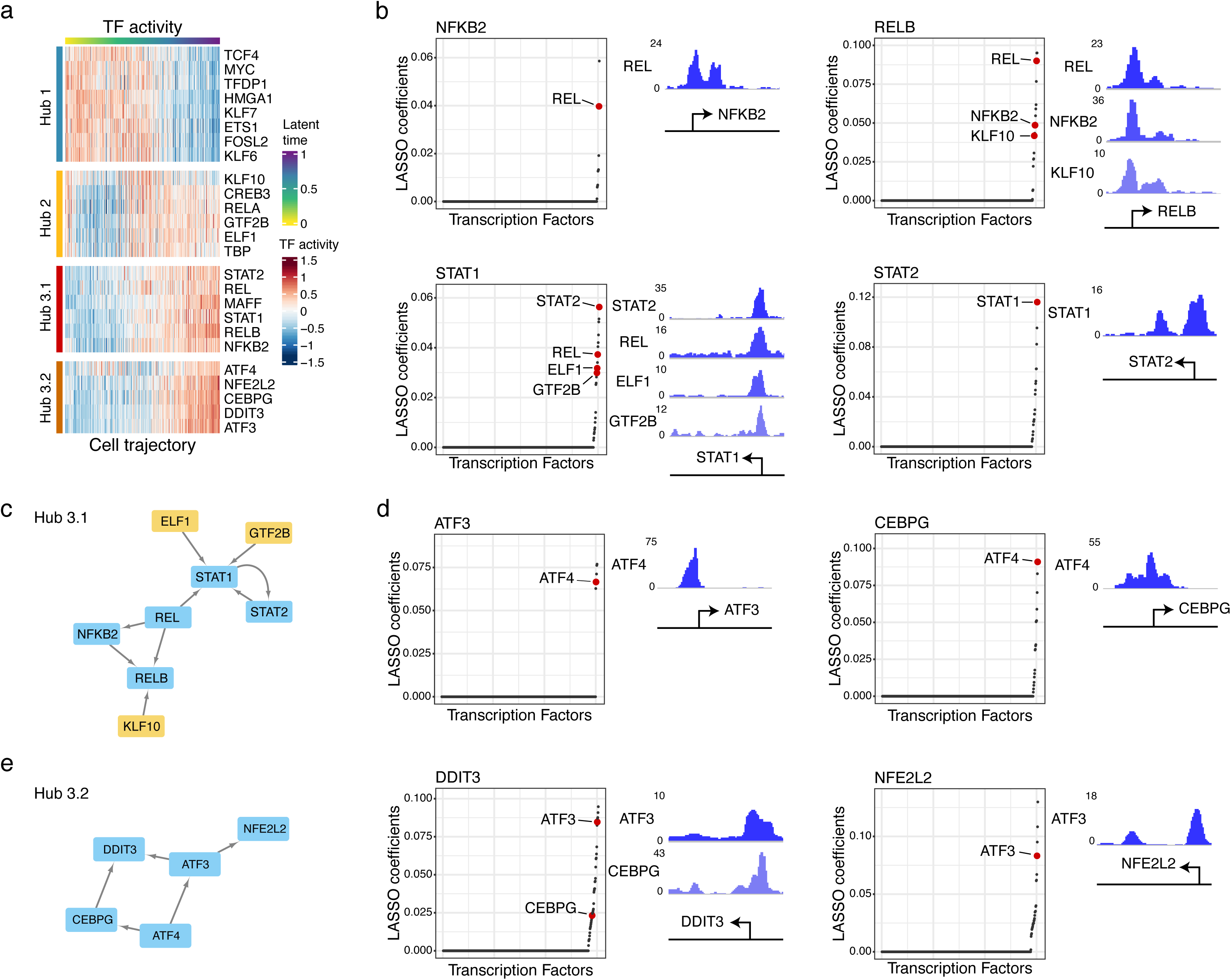
Identifying key TF regulatory networks driving OIS. (**a**) Heatmap of the activities of the verified TFs. (**b**) Establish the regulatory links between the TFs within TF hub 3.1 by correlating the changes in the nascent RNA of the TF of interest with the changes in the mature RNA of the other TFs. The direct regulatory links (colored in red) were identified by ChIP-seq verification. The corresponding ChIP-seq peaks were plotted on the right. (**c**) The TF regulatory network of TF hub 3.1. The regulatory network was established based on the direct regulatory links between different TFs. The TFs belonging to TF hub 2 are colored in yellow. (**d**) Establish the regulatory links between the TFs within TF hub 3.2 by correlating the changes in the nascent RNA of the TF of interest with the changes in the mature RNA of the other TFs. The direct regulatory links (colored in red) were identified by ChIP-seq verification. The corresponding ChIP-seq peaks were plotted on the right. (**e**) The TF regulatory network of TF hub 3.2. The regulatory network was established based on the direct regulatory links between different TFs.

Notably, based on the gene expression kinetics of the 25 TFs, Hub 3 can be further divided into two sub-hubs (Hub 3.1 and Hub 3.2). Intriguingly, Hub 3.1 predominantly consisted of TFs regulating the immune response pathway, while Hub 3.2 was comprised of TFs linked to the integrated stress response pathway (**Fig. 6a**). This observation highlights the distinct and specific functional roles of different TF hubs during the entry into the OIS.

In Hub 3.1, we identified *REL* as a critical player that orchestrates the activities of both the NF-kB pathway and the STAT1/STAT2 pathway (**Fig. 6b-c**). Interestingly, we also observed that the TFs belonging to Hub 2 (colored in yellow in **Fig. 6c**) were also involved in the regulation of *RELB* and *STAT1* through the direct binding to their promoter regions, confirming the previous observation of the regulatory connection between Hub 2 and Hub 3.

In Hub 3.2, *ATF4* was identified as the most upstream regulator within this regulatory network. Our data revealed that *ATF4* directly regulated the expression of *CEBPG* and *ATF3* (**Fig. 6d-e**), both of which play critical roles in regulating cellular stress response ^50,51^. The activation of *CEBPG* and *ATF3*, in turn, led to the upregulation of *DDIT3,* which is involved in the regulation of ER stress response^52,53^, and *NFE2L2*, which regulates the oxidative stress response^54^ (**Fig. 6d-e**). This cascade of regulatory events underscores ATF4’s central role in orchestrating the stress response pathway through the regulation of key downstream genes, aligning with previous findings^55–58^. Overall, our data unveiled the core TF network that governs the entry into cellular senescence after the induction of oncogenic stress.

### RNA binding proteins (RBP) and alternative polyadenylation (APA) based post-transcriptional regulations during OIS

As shown in **Fig. 5e**, a substantial proportion of genes were Type II DEGs, whose gene expression is dominantly affected by post-transcriptional regulation. Based on the dynamic changes in their mature RNA abundance, we classified these genes into five kinetic modules with significant enrichment of different pathways (**Supplementary** Fig. 10). Next, we examined the potential mechanisms driving the differential gene expression of Type II DEGs. We first compared the binding targets of the known 150 RBPs with these five kinetic modules, and we identified the significant enrichment of the binding targets of certain RBPs in Modules 1, 2, and 3 (**Supplementary Fig.11a**). Interestingly, a notable proportion of the RBPs enriched in Module 1 and 2 are splicing regulators^39^ (**Supplementary Fig.11b**), indicating that the splicing of the target genes detected in these two modules was regulated during oncogene-induced senescence.

Besides the regulation of RNA fate by RBPs, alternative polyadenylation (APA) has also been reported as a common mechanism to regulate gene expression in response to cellular stress^40^. The lengthening of 3’UTR associated with the usage of distal polyA sites could increase the binding sites for RBPs or miRNAs and hence promote RNA degradation^59–61^. Interestingly, Chen *et al.* have reported a global lengthening of 3’UTR in replicative senescence^62^. To test whether APA is also involved in the post-transcriptional regulation in OIS, we evenly separated the cells into five time windows (intervals) along the trajectory and used DaPars2^63^ to derive the preference of polyA site usage within each interval. Indeed, a substantial proportion of the genes were found to use multiple polyA sites (**Supplementary** Fig. 12a), which makes them qualified for APA analysis.

As a result, we observed different trends in alternative polyadenylation among the five kinetic modules. In Module 1, we observed an increase in the usage of distal polyA sites in interval 2 and interval 5, which corresponds to the decrease in gene expression observed at the corresponding intervals (**Supplementary** Fig. 12b-c). In Module 3, 4 and 5, we observed a clear decrease in the usage of distal polyA sites at the late intervals (**Supplementary** Fig. 12g**-i**), which is consistent with the significant increase in the expression levels of these genes at the late time points (**Supplementary** Fig. 12f). It is worth noting that in Module 2, we only observed slight changes in the usage of polyA sites along the trajectory (**Supplementary** Fig. 12d-e), suggesting that the expression of these genes was not mainly regulated by APA. Overall, these observations confirm APA as a general post-transcriptional regulation during oncogene-induced senescence.

## Discussion

In summary, we developed a high-sensitivity single-cell total-RNA-seq method (snapTotal-seq) by combining the multiple annealing chemistry and template switching chemistry into a one-step reaction. By taking the intrinsic cell cycle as an example, we conducted a comprehensive benchmark analysis to compare both oligo-dT based and total-RNA based scRNA-seq methods for their performance in characterizing gene expression dynamics through RNA velocity analysis. As a result, with the efficient detection of both nascent RNAs and mature RNAs, snapTotal-seq achieved the best confidence score in the gene expression kinetic analysis.

With the established cell trajectory by RNA velocity, we identified a large number of novel cell-cycle genes and unveiled the related biological processes intricated with cell-cycle progression. More importantly, by LASSO regression of the mature RNA expression levels of TFs against the transcriptional dynamics of the gene of interest represented by nascent RNAs, we were able to directly link the expression of TFs to the transcriptional activities of their target genes. In contrast, with methods that only profile the mature RNAs, this analysis is unfeasible. It is also worth noting that even for the methods using nucleotide analogs that label newly synthesized RNA, these labeled RNAs are also based on mature RNAs. Therefore, it does not directly capture the transcription activities represented by nascent RNAs.

To zoom into the regulatory network of TFs, we can perform LASSO regression of the mature RNA expression levels of a specific TF against the transcriptional dynamics of other TFs. As a result, we demonstrated the ability of snapTotal-seq to derive the inference hubs of TFs and identify the key TFs that control the cell-state transition. To further illustrate the performance of snapTotal-seq, we applied it to the process of oncogene-induced senescence. And we successfully pinpointed the key TF regulatory hubs that drove the cells into the senescent state.

In summary, our work demonstrates the versatility and potential of snapTotal-seq in advancing our understanding of gene expression dynamics and, more importantly, the regulatory hubs underlying cell-state transition.

## Author contributions

C.Z. and Y.N. designed the project. Y.N. and J.L. performed the experiments. Y.N. and C.Z. performed data analysis. C.Z. and Y.N. wrote the manuscript.

## Acknowledgments

We are grateful to the McNair family for their support. We thank the Cytometry and Cell Sorting Core at Baylor College of Medicine for their kind help on the FACS-related experiments. We would also like to thank other Zong lab members for their kind support. This work was supported by the McNair Scholarship.

## METHODS

### Cell Cultures

HEK293T cells were cultured in DMEM with 10% fetal bovine serum (FBS, Life Technologies) and were passaged every two days with 0.05% trypsin (Corning^®^). HPNE cell line was cultured in the medium with 75% DMEM without glucose (Sigma), 25% Medium M3 Base (INCELL Corp.), 2mM L-glutamine (Sigma), 1.5 g/L sodium bicarbonate (Sigma), 5% FBS (Life Technologies), 10 ng/mL human recombinant EGF (ThermoFisher), 5.5mM D-glucose (Sigma) and 750 ng/mL puromycin (InvivoGen), as recommended by ATCC. The cells were passaged every 2-3 days with 0.25% trypsin (Corning^®^).

The induction of oncogene-induced senescence was performed as described in previous publicaiton^49^. Briefly, wild-type HPNE cells with inducible KRAS^G12D^ (iKRAS-HPNE) were cultured with doxycycline (6 μg/mL) to activate the expression of KRAS^G12D^. The cells were collected on day 1, day 2, day 3, and day 5 after the induction of KRAS^G12D^ expression for single-cell RNA-seq experiments.

U2OS cell line was cultured in McCoy’s 5a medium (ATCC) supplemented with 10% FBS (Life Technologies), and the cells were passaged every 2-3 days with 0.25% trypsin (Corning^®^). NIH/3T3 cell line was cultured in DMEM with 10% fetal bovine serum (FBS, Life Technologies) and was passed every 2 days with 0.25% trypsin (Corning^®^). To perform single-cell RNA-seq experiments, the cells were trypsinized and resuspended in PBS (Corning^®^). The cells were then sorted into the 96-well plates with 1 μL lysis buffer per well by using BD Aria II with a 130 μm nozzle.

### Cell lysis, reverse transcription, and amplification by snapTotal-seq

The cell lysis buffer consisted of 0.7 μL of 1.8% Triton-X (Sigma), 0.025 μL of 0.1 M DTT (Invitrogen), 1 U RNaseOUT (Invitrogen), 0.05 μL of dNTP (10 mM each) and 0.2 μL of primer mix (1.5 μM of GTG AGT GAT GGT TGA GGA TGT GTG GAG N5 T12, 5 μM of GTG AGT GAT GGT TGA GGA TGT GTG GAG N5 T3, 5 μM of GTG AGT GAT GGT TGA GGA TGT GTG GAG N5 G3). 10 μL of mineral oil (Sigma) was added to prevent the evaporation in the following steps. The lysis was performed at 42 °C for 3.5 minutes. After lysis, the plate was immediately placed on the ice for 1 minute. Reverse transcription mix which contained 0.4 μL of 5X M-MLV reverse transcriptase buffer (250 mM Tris-HCl, 375 mM KCl, 15 mM MgCl_2_, Invitrogen), 0.1 μL of 0.1 M DTT (Invitrogen), 2 U RNaseOUT (Invitrogen), 10 U Maxima H Minus Reverse Transcriptase (Thermo Scientific) and 0.4 μL of 0.1% Triton-X (Sigma), was then added to each well. The reverse transcription and template switching step was carried out with 10 cycles of 8 °C for 12 s, 15 °C for 45 s, 20 °C for 45 s, 30 °C for 30 s, 42 °C for 2 min, and 50 °C for 3 min, followed by 50 °C for 15 min. The reverse transcriptase was inactivated at by incubating at 74 °C for 25 min.

After that, PCR amplification mix, which consisted of 10 μL of 5X GoTaq Flexi buffer, 4 μL of 25 mM MgCl_2_, 1 μL of dNTP (10 mM each), 0.25 μL of 100 μM GAT primer (GTG AGT GAT GGT TGA GGA TGT GTG GAG), 1.75 U GoTaq Flexi DNA polymerase, 2.5 μL of 20X EvaGreen Dye (Biotium) and 29.9 μL of RNase-free H_2_O, was added to each well. The amplification was carried out on a Real-time PCR machine. The PCR program was as follows: 95 °C for 2 min, 23-26 cycles of 95 °C for 20 s, 63 °C for 20 s and 72 °C for 2 min, 72 °C for 5 min.

Purification was carried out with 1.2X Ampure XP beads (Beckman Coulter). The samples were mixed with Ampure XP beads and incubated for 10 min at room temperature. Then, the plate was placed on a 96-well magnetic stand, and the supernatant was removed. To remove the residual mineral oil, we washed the beads with 2-propanol (Sigma) twice, followed by the wash with 100% ethanol (Koptec). Next, the beads were washed twice with 80% ethanol. Finally, the amplified products were eluted in 25 μL of RNase-free H_2_O.

### Library construction

Firstly, the amplified products from each cell were tagged with cell-specific barcodes by double-strand conversion (DSC). Specifically, for each cell, 10 μL of amplified product was mixed with 2 μL 10X ThermoPol Buffer, 0.4 μL of dNTP (10mM each), 1 μL of 10 μM DSC primer with cell barcode, 0.4 U Deep Vent (exo-) DNA polymerase (New England BioLabs) and RNase-free H_2_O to 20 μL. The DSC program was as follows: 95 °C for 1 min, 15 cycles of 63 °C for 25 s and 72 °C for 1 min, 72 °C for 3 min. 1 μL of 0.2 M EDTA (Sigma) was then added to each cell to stop the reaction.

The cells with different cell barcodes were pooled together (4 μL of reaction product per cell) and were purified with 1.1X Ampure XP beads. For each pooled library, ∼350 ng of purified DNA products were then sonicated to 150-250 bp (Covaris S220). The sonicated DNA was purified with 1.8X Ampure XP beads. Following that, another step of DSC was performed to enrich the DNA fragments with UMI and cell barcode. Briefly, the sonicated product was mixed with 3 μL of 10X ThermoPol Buffer, 0.6 μL of dNTP (10 mM each), 1.5 μL of 10 μM primer (GCACGACATCTGCTAACGCAGTA GTGTGCTCTTCCGATCT), 0.6 U Deep Vent (exo-) DNA polymerase and H_2_O to 30 μL. The reaction was carried out as follows: 95 °C for 1 min, 6 cycles of 56 °C 25 s and 72 °C 30 s, 72 °C for 3 min. The products were then purified with 1.4X Ampure XP beads. The purified products were subjected to dA tailing with 2 μL of 10X NEBuffer 2 (New England BioLabs), 0.1 μL of 100 mM dATP, and 2.5 U Klenow Fragment (3’ -> 5’ exo-) (New England BioLabs) and H_2_O to 20 μL, by incubating at room temperature for 30 min.

After purifying the products with 1.4X Ampure XP beads, we performed the ligation at room temperature for 20 min. The ligation reaction mix included 13 μL of 2X Quick Ligase reaction buffer, 0.5 μL of 50 mM Y-shape adapter, 0.5 μL of Quick Ligase (New England BioLabs), and 12 μL of dA-tailing product. The ligation reaction was quenched by adding 5 μL of 0.2 M EDTA (Sigma) and was purified by 1.2X Ampure XP beads. The amplification of the ligation products was then performed with the program as follows: 95 °C for 2 min, 10 cycles of 95 °C 20 s, 61 °C 20 s and 72 °C for 1 min, and 72 °C for 3 min for final extension. The amplification mix consisted of 5 μL of ThermoPol Buffer, 1 μL of dNTP (10 mM each), 2 μL of 10 μM forward primers (GCA CGA CAT CTG CTA ACG CAG TA), 2 μL of 10 μM reverse primers (AAT GAT ACG GCG ACC ACC GAG A), 1 U of Deep Vent (exo-) DNA polymerase, 33.5 μL of H_2_O and 6 μL of ligation products.

After the amplification products were purified with 1.2X Ampure XP beads, duplex-specific nuclease (DSN) treatment was applied to remove ribosomal reads. Specifically, 100 ng of amplified products was mixed with 2 μL of 10X DSN buffer (Evrogen) and H_2_O to 20 μL. The DNA was denatured at 95 °C for 30 s, followed by incubation at 80°C for 3 hours. After that, 1 μL of preheated duplex-specific nuclease (Evrogen) was added to the reaction and incubated at 80 °C for 15 min. To quench the reaction, 4 μL of 0.2 M EDTA was added at 80 °C, and the reaction was then put on ice immediately. The products were purified with 1.2X Ampure XP beads.

Following that, an enrichment PCR was carried out to enrich the undigested DNA fragments. The reaction mix consisted of 2.5 μL of 10X ThermoPol Buffer, 0.5 μL of dNTP (10 mM each), 0.75 μL of 10 μM forward primers (GCA CGA CAT CTG CTA ACG CAG TAG TGT GCT CTT CCG ATC T), 0.75 μL of 10 μM reverse primer (AAT GAT ACG GCG ACC ACC GAG A), 0.5 U Deep Vent (exo-) DNA polymerase, 12.25 μL of H_2_O and 8 μL of DSN treatment product. The program was as follows: 95 °C for 2 min, 5 cycles of 95 °C for 20 s, 63 °C for 20 s and 72 °C 1 min, 72 °C 3 min for final extension. The amplified products were purified with 1.2X Ampure XP beads.

A final PCR step, which consisted of 2.5 μL of 10X ThermoPol Buffer, 0.5 μL of dNTP (10 mM each), 0.75 μL of 10 μM forward primers (CAA GCA GAA GAC GGC ATA CGA GAT GCA CGA CAT CTG CTA ACG CAG TA), 0.75 μL of 10 μM reverse primers (AAT GAT ACG GCG ACC ACC GAG A), 0.5 U Deep Vent (exo-) DNA polymerase, 19.25 μL of H_2_O and 1 μL of purified DNA product, was performed to add the sequencing adapter. The program runs as follows: 95 °C for 2 min, 5 cycles of 95 °C for 20 s, 61 °C for 20 s and 72 °C 1 min, 72 °C 3 min for final extension.

The libraries were sequenced on NextSeq 500 machine with customized sequencing primers as follows: Read 1 sequencing primer: ACA CTC TTT CCC TAC ACG ACG CTC TTC CGA TCT (the same as Illumina Tru-seq i5 sequencing primer), Read 2 sequencing primer: AGA GGT GAG TGA GTG ATG GTT GAG GAT GTG TGG AG, Index i5 sequencing primer: AGA TCG GAA GAG CGT CGT GTA GGG AAA GAG TGT, Index i7 sequencing primer: CTC CAC ACA TCC TCA ACC ATC ACT CAC TCA CCT CT. Read 1 sequenced the captured RNA sequence, while read 2 sequenced the UMI.

### Reads alignment and gene expression calculation

Read 1 was mapped to human genome assembly (GRCh37) by using STAR (v2.5.3a)^64^. The uniquely mapped reads were then mapped to the gene annotations of GENCODE (v19) by using htseq-count^65^ with ‘intersection-strict’ mode and with option ‘--stranded=no’. To discern the amplicons from exons and introns, the ‘transcript’ feature and the ‘exon’ feature were used respectively. The UMI sequences (the first five bases of read 2) of the reads mapped to the gene regions (either the ‘transcript’ feature or the ‘exon’ feature) were extracted. The reads were then grouped by the UMI sequence and the gene that they were mapped to, as these reads were derived from the same original cDNA amplicon. If all the reads within the group were mapped to the exon regions of the corresponding gene, the original amplicon was classified as an exonic amplicon. Otherwise, the original amplicon was classified as an intronic amplicon. The number of exonic amplicons and intronic amplicons were then counted for each gene, and the exonic UMI count matrix and intronic UMI count matrix were generated.

### Normalization, PCA, cell cycle analysis, and RNA velocity analysis

The genes that were detected with more than 1 UMI in at least five cells in exon data were kept. The rest of the genes were defined as lowly expressed genes and were discarded in the following analysis. The mitochondria genes were also removed before the normalization step. To normalize the gene expression data across different cells, we divided the count of UMIs of each gene by the total UMIs detected in each cell and multiplied by the average UMI number of all cells. The normalization step was performed on exonic data and intronic data separately. PCA was performed on exon-based gene expression data by using Seurat package^66^. Briefly, the exonic normalized gene expression data were log-transformed and scaled. The top 500 most variable genes were selected by using ‘FindVariableFeatures’ function to perform PCA. The cell cycle analysis was performed with reCAT^22^ by using the exon-based gene expression data. The log-transformed normalized gene expression was used as the input. RNA velocity analysis was performed by using scVelo package^17^. The raw count matrices were used as the input. To filter the lowly expressed genes, the ‘min_shared_counts’ was set as 30 and the ‘min_cells_u’ as 5. After normalization and log-transformation, the velocities were projected by using the top 2000 most variable genes and the ‘dynamical modeling’ mode with the following parameters: n_pcs=2, n_neighbors=20, fit_basal_transcription=True. Latent time was calculated based on the projected RNA velocities.

### Cell cycle trajectory based differential gene expression analysis

The differential gene expression analysis was performed on exonic data and intronic data, respectively, by using tradeSeq package^23^. The exonic or intronic raw count matrix was used as the input. The pseudotime of each cell was derived from the cell cycle trajectory inferred by RNA velocity. The parameter ‘nknots’ in function ‘fitGAM’ was set as 5. The ‘associationTest’ function was used to identify the genes which were differentially expressed along the pseudotime. The genes whose exonic normalized gene expression values and intronic normalized gene expression values were ≥ 3 in at least 10 cells were kept. To perform the multiple test correction, the false discovery rate (FDR) was calculated by using p.adjust function in R. To verify the results of DEG analysis, we calculated the fold-changes in gene expression along the cell cycle for all genes. To account for the variations in single-cell data, we evenly separated the cells into 6 intervals along the cell cycle, and the average expression levels were calculated for each interval. The fold changes between the highest expression level and the lowest expression level among these intervals were then calculated.

### Benchmark analysis on Smart-seq3, CEL-Seq2, VASA-seq and Smart-seq-total

To perform the benchmark analysis on Smart-seq3, we reanalyzed the published HEK293FT dataset generated by Smart-seq3. The raw fastq files were first demultiplexed based on the cell indexes. The UMI reads were identified based on the adaptor sequence. The adaptor sequence and the UMI sequence were first trimmed by using seqtk. The trimmed reads were then mapped to human genome assembly (GRCh37) by using STAR (v2.5.3a). Following that, the uniquely mapped reads were assigned to exon or intron regions by using htseq-count as described above. The reads were collapsed if the hamming distance of UMIs ≤ 1, and the exonic UMI count matrix and intronic UMI count matrix were generated. Two potential outliers that were identified based on the PCA plot in the initial analysis were discarded. The normalization, PCA, cell cycle analysis, and RNA velocity analysis were then performed as described above.

To perform the benchmark analysis on CEL-Seq2, we reanalyzed the published HEK293 dataset generated by CEL-Seq2. To avoid potential batch effects, only the cells collected in the mixture 2 experiment, as described in the original study, were used. The demultiplexed reads were mapped to human genome assembly (GRCh37) by using STAR (v2.5.3a). Next, the uniquely mapped reads were assigned to exons or introns, as described above. The reads that were mapped to the same gene were collapsed based on their UMI sequences, which generated the exonic UMI count matrix and intronic UMI count matrix. The potential outliers that were identified based on the PCA plot in the initial analysis were discarded. The normalization, PCA, cell cycle analysis, and RNA velocity analysis were then performed as described above.

To reanalyze the published HEK293T dataset generated by VASA-seq (plate version), the raw fastq files were first demultiplexed based on the cell indexes. The demultiplexed reads were mapped to human genome assembly (GRCh37) by using STAR (v2.5.3a). Next, the uniquely mapped reads were assigned to exons or introns, as described above. The reads that were mapped to the same gene were collapsed based on their UMI sequences, which generated the exonic UMI count matrix and intronic UMI count matrix. The cells with low sequencing depth (total exon UMI < 7,500) or high genomic DNA contamination were discarded. The potential outliers which were identified based on the initial PCA were also discarded. The normalization, PCA, cell cycle analysis, and RNA velocity analysis were then performed as described above.

To analyze the published HEK293T dataset generated by Smart-seq-total, we first trimmed the poly(A) sequences from the raw reads using Cutadapt (v3.4)^67^. The trimmed reads were then mapped to human genome assembly (GRCh37) by using STAR (v2.5.3a). The uniquely mapped reads were assigned to exon or intron regions by using htseq-count as described above.

### Validation of the identified cell cycle dependent genes by qRT-PCR

HEK293T-FUCCI cell line was established by using the FastFUCCI plasmid (addgene #86849). The cells with successful transduction were selected by culturing the cells with puromycin (1 μg/mL) for four days. After collecting the cells of different cell cycle phases by FACS, RNA was extracted by using TRIzol reagent (Invitrogen) according to the manufacturer’s instructions. The reverse transcription reaction and qPCR were then carried out by using iScript™ reverse transcription supermix (Bio-Rad) and iTaq universal SYBR green supermix (Bio-Rad), respectively. The qPCR program was as follows: 94 °C for 2 min, 40 cycles of 94 °C for 20 s, 58 °C for 20 s, and 72 °C for 20 s. *ATP5F1* was used as the internal control, as its expression remained unchanged along the cell cycle.

### Calculation of cross-correlation

To quantify the coupling between the gene expression changes in nascent RNA and mature RNA, we borrowed the metrics of ‘cross-correlation’ from the field of signal processing, which is defined as a measure of similarity between two signals. For each gene module, the average z-score of gene expression along the cell cycle trajectory was calculated for nascent RNA and mature RNA, respectively. Following that, the cross-correlation between the expression curves of nascent RNA and mature RNA was calculated by using numpy.correlate function in Python.

### Functional analysis on cell cycle genes (CCGs)

The list of known CCGs was obtained by combining the gene list of the cell cycle pathway in the Gene Ontology database^68^, the gene list of the cell cycle pathway in the Reactome database^69^ and the genes reported in Cyclebase^70^. The protein-protein interaction (PPI) analysis was carried out by using STRING database^71^ with the default parameters. Cytoscape (v3.8.2) was used for the PPI network visualization. The Gene Ontology enrichment analysis was performed by using ‘Hypergeometric’ model in ‘goseq’ R package^72^. The cellular localization was obtained from the cellular compartment category of the GO database. Fisher’s exact test was used to determine the differential enrichment of known CCGs and novel CCGs in different cellular compartments. The ‘endomembrane system’ consisted of ‘GO:0031226’, ‘GO:0005887’, ‘GO:0031224’, ‘GO:0016021’, ‘GO:0000139’, ‘GO:0098791’, ‘GO:0098588’, ‘GO:0005783’, ‘GO:0005794’, ‘GO:0031090’, ‘GO:0031982’ and ‘GO:0012505’. The list of genes localized in the nucleus was obtained from ‘GO:0005634’.

### Identify the regulatory links between TFs and their target genes

The links between TFs and their target genes were first identified based on co-expression analysis. The list of annotated TFs was obtained from RcisTarget package^26^ and human TF database^73^. Only the TFs annotated in both databases were selected. The gene expression at the mature RNA and nascent RNA levels were calculated as described above. The normalized expression values were log-transformed, centered, and scaled. For each gene, the correlation coefficients between its expression at the nascent RNA level and the expression of all TFs at the mature RNA level were calculated. The TFs with r > 0.15 were selected to construct the LASSO regression model with glmnet package. The LASSO regression model was constructed to predict the nascent RNA expression of the gene of interest based on the expression of selected TFs. As a negative correlation could be caused by the mutual exclusive gene expression patterns, only a positive correlation was considered here. The links with regression coefficients > 0.03 were considered. The linked genes of each TF were then compared to Type I kinetic modules. The significant enrichment was determined by FDR < 0.01, enrichment fold > 2, and the number of overlapped genes > 10. The enriched TF-gene links were then subjected to ChIP-seq verification or motif enrichment analysis

### Establishing TF regulatory network

The links between different TFs were identified based on the expression covariance as described above. All of the identified links were then verified by published ChIP-seq data. The verified TF-TF links were used to establish the TF regulatory network with Cytoscape^74^. For the TF association network in Fig. 5j, all TF-TF links were used.

### ChIP-seq analysis and motif enrichment analysis

The peak files were downloaded from CistromeDB^25^. The samples with median mapping quality ≥ 25, unique mapping rate ≥ 0.6, PCR bottleneck coefficient ≥ 0.8, a fraction of reads in peaks ≥ 0.01, peak number (fold > 10) ≥ 150, and high consistency with DNase-seq data (percentage of top 5k peaks overlapped with DNase-seq ≥ 0.85) were kept. The rest of the datasets were not used due to potentially low quality. The qualified datasets were then grouped based on their target TFs. For each TF, the peaks (fold > 10) identified in corresponding datasets were merged, and the peak annotation was performed using HOMER^75^. The binding targets were identified if the peak was within 1kb or 5kb from the transcription start site. These two different cutoffs corresponded to the scenarios of binding to the promoter region or binding to the enhancer region, respectively.

For ChIP-seq peak visualization, the raw data or the processed data were downloaded. The processed data were directly used for visualization in Integrative Genomics Viewer (IGV)^76^. Raw data were mapped to human genome assembly (GRCh37) using Bowtie2^77^ after removing the adapter sequences with Cutadapt. After deduplication, BAM files were converted to bigwig files using bamCoverage function in deepTools package^78^. Bigwig files were then used for visualization in IGV. The following datasets were used for visualization: E2F1, GSM2132552; E2F2, ENCFF826PYA; KLF11, GSE59703; MYBL2, ENCFF487KQK; REL, GSE55105; NFKB2, GSE55105; KLF10, ENCFF437PEU; STAT2, ENCFF133YDB; ELF1, ENCFF937HON; GTF2B, GSE71848; STAT1, GSE43036; ATF4, GSE69309; ATF3, ENCFF783IOJ; CEBPG, ENCFF182SSK.

Motif enrichment analysis was carried out within 500bp upstream of TSS or 5kb around the TSS using RcisTarget package^26^. The significant enrichment in motif analysis was determined by normalized enrichment score (NES) > 3 and the number of enriched genes ≥ 10.

### Gene knockdown with CRISPR interference (CRISPRi)

HEK293T-CRISPRi cell line was established by using dCas9-KRAB-MeCP2 plasmid (addgene #122205). The cells that were successfully transduced were selected with 10 μg/mL Blasticidin (InvivoGen) for a week. The sgRNA vector was generated based on CROPseq-Guide-Puro (addgene #86708). To enable the measurement of the cell population transduced with sgRNA vector via fluorescence, we replaced the puromycin resistance gene (PuroR) with RFP. Briefly, the CROPseq-Guide-Puro vector was digested by using MluI (NEB) and SmaI (NEB), which was then ligated with the RFP sequence from pLKO5.sgRNA.EFS.tRFP plasmid (addgene #57823). To clone the sgRNA sequence into the sgRNA vector, the sgRNA vector was digested by using Esp3I (ThermoFisher), and the digested vector was next ligated with the annealed sgRNA oligo by using T4 ligase (New England BioLabs). The lentivirus was prepared, and the sgRNA was then transduced into the HEK293T-CRISPRi cell line. After three days of transduction, the transduced cells were 1:1 mixed with uninfected HEK293T-CRISPRi cells. The rest of the cells were harvested for RNA extraction, and qRT-PCR was performed to verify the knockdown efficiency by using *ACTB* as the internal control. After five days of transduction, the initial percentage (referred to as day 0) of RFP^+^ cells in the mixed cell population was measured by using flow cytometry. The mixed cell population was further cultured for 14 days, and the percentage of RFP^+^ cells was measured by using flow cytometry after 14 days of culturing to determine the effects of the target gene on cell proliferation. The relative proliferation was calculated as RFP^+^% (day 14) / RFP^+^% (day 0).

### Data analysis of the gene expression dynamics during oncogene-induced senescence

After mapping and UMI counting, lowly expressed genes were filtered out, and normalization was performed as described above. PCA was carried out with the top 1000 most variable genes. The first five principal components were selected to generate UMAP. Cell cycle analysis was performed on the cells collected at each time point respectively by using reCAT. RNA velocity analysis was performed by using the scVelo package. The raw count matrices were used as the input. To filter the lowly expressed genes, the ‘min_shared_counts’ was set as 50 and the ‘min_cells_u’ as 10. After normalization and log-transformation, the velocities were projected by using the top 2000 most variable genes and the ‘dynamical modeling’ mode with the following parameters: n_pcs=5, n_neighbors=20. G0 cells were then ordered based on the latent time to establish the trajectory toward oncogene-induced senescence. The differential gene expression analysis was performed based on the established trajectory by using tradeSeq as described above. The Gene Ontology enrichment analysis was performed on each kinetic module by using ‘Hypergeometric’ model in ‘goseq’ R package.

### Data availability

The raw data of Smart-seq3 were obtained from ArrayExpress: E-MTAB-8735 at EMBL-EBI. The raw data of CEL-Seq2 were obtained from the Gene Expression Omnibus: GSE132044 (SRR9167461, SRR9167462, SRR9171229, SRR9171230). The raw data of the VASA plate were obtained from the Gene Expression Omnibus: GSE176588. The raw data of Smart-seq-total were obtained from the Gene Expression Omnibus: GSE151334. The raw data and the processed data sets generated in this study are available at the Gene Expression Omnibus under the accession number GSE202126.

### Code availability

The analysis pipeline customized for snapTotal-seq sequencing data is available at https://github.com/zonglab/snapTotal-seq.git.

