## Supplementary Materials for "Single-cell Total-RNA Profiling Unveils Regulatory Hubs of Transcription Factors"

# Of

[4]Integrative Molecular and Biomedical Sciences Program

[5]Dan L Duncan Comprehensive Cancer Center,

[6]McNair Medical Institute,

Baylor College of Medicine,

One Baylor Plaza, Houston, Texas, 77030

[*]These authors contributed equally to this work

a

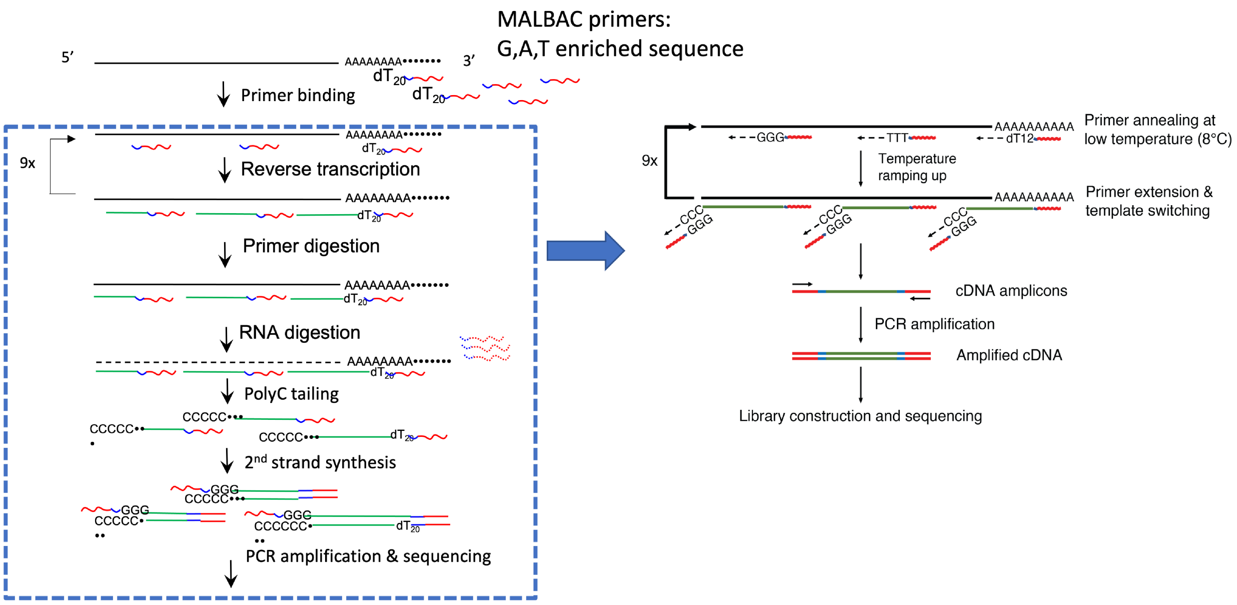


b

| Method | RNA fragmentation | poly(A) tailing | Reverse transcription | Second-strand synthesis | Amplification |
|---|---|---|---|---|---|
| snapTotal-seq | / | / | Yes (with TS) | / | PCR |
| VASA-seq | Yes | Yes | Yes | Yes | IVT |
| Smart-seq-total | / | Yes | Yes (with TS) | / | PCR |
| Smart-seq3 | / | / | Yes (with TS) | / | PCR |
| CEL-Seq2 | / | / | Yes | Yes | IVT |

**Supplementary figure 1.** Chemistry of snapTotal-seq. (**a**) The comparison between the chemistries of MATQ-seq (left) and snapTotal-seq (right). (**b**) Summary of the major steps in snapTotal-seq, VASA-seq, Smart-seq-total, Smart-seq3 and CEL-Seq2. TS, template switching. IVT, in vitro transcription.

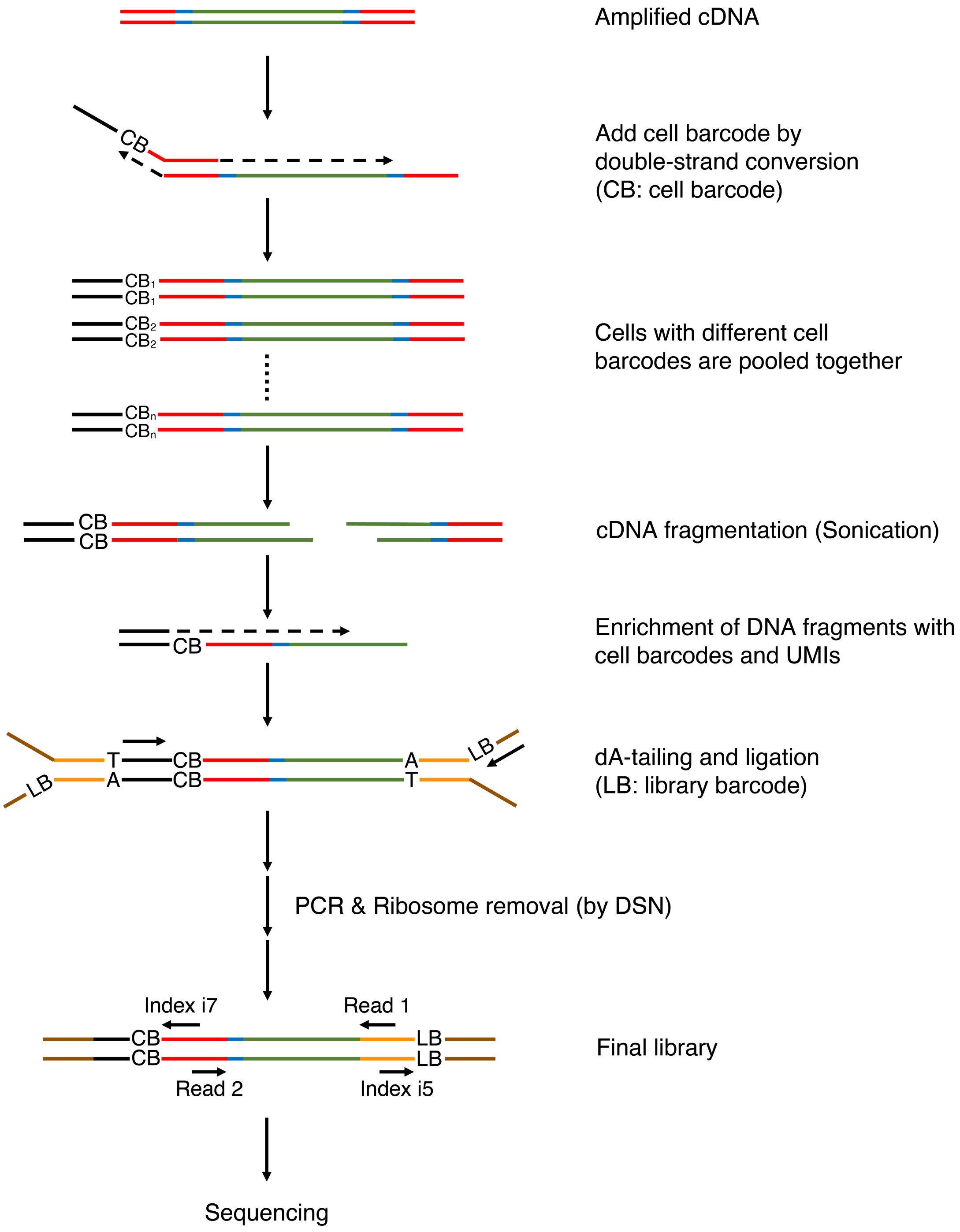


**Supplementary figure 2.** Scheme of library construction.

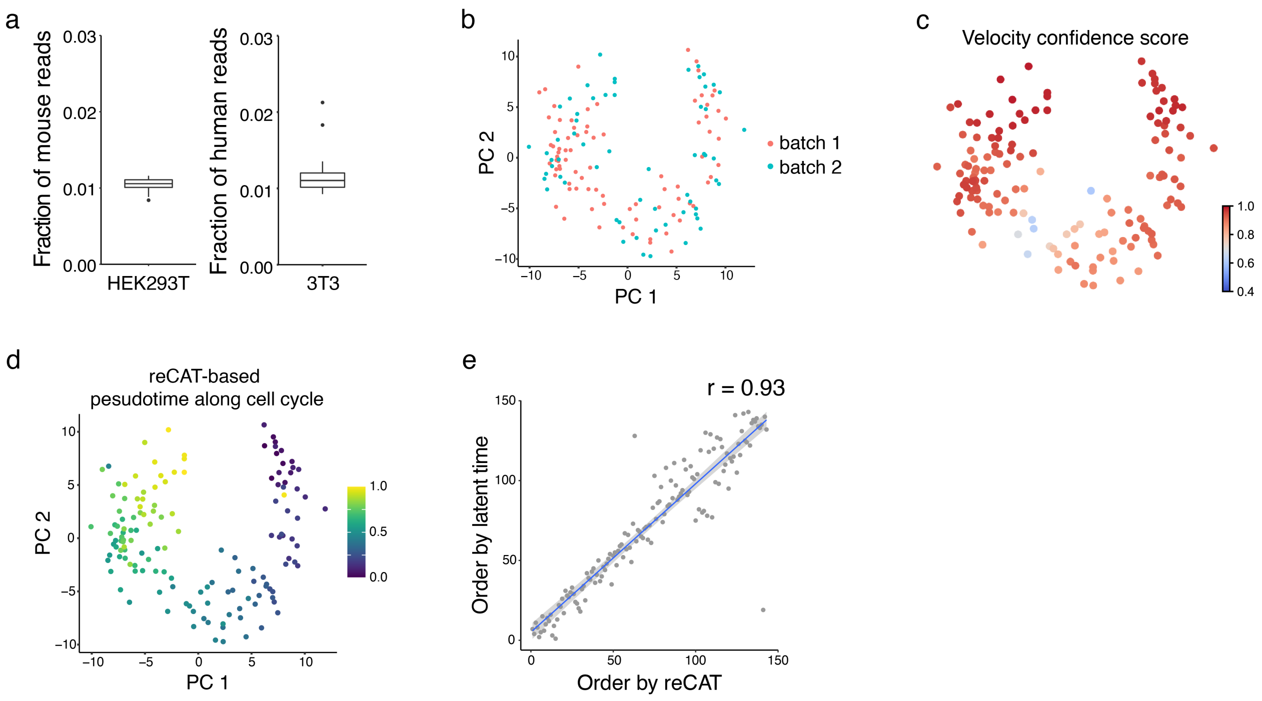


**Supplementary figure 3.** Benchmark analysis. (**a**) Fraction of reads from the other species. (**b**) PCA plot of HEK293T cells. The cells are colored by the corresponding technical batches. (**c**) Velocity confidence scores of HEK293T cells sequenced by snapTotal-seq. (**d**) The pseudo-temporal trajectory along cell cycle established by reCAT algorithm. (**e**) The scatter plot between the reCAT based cell cycle trajectory and the RNA velocity based cell cycle trajectory.

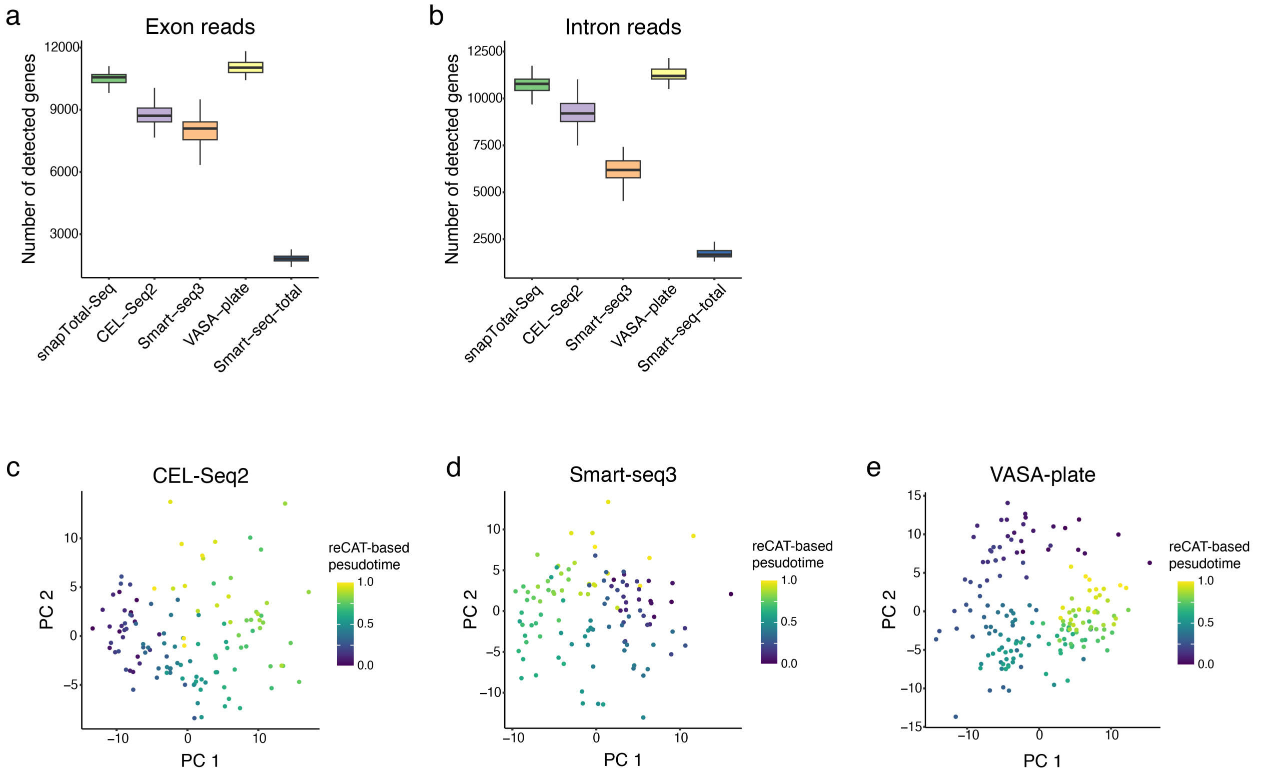


**Supplementary figure 4.** Benchmark analysis. (**a-b**) The number of genes detected by exon reads or intron reads in single HEK293T cells by different methods. Panel **a**, exon reads. Panel **b**, intron reads. For equal-footing comparison, all tested cells were downsampled to the depth of 1M uniquely mapped reads per cell. (**c-e**) The cell cycle pseudotime inferred by using reCAT for CEL-Seq2 (**c**), Smart-seq3 (**d**) and VASA-plate (**e**).

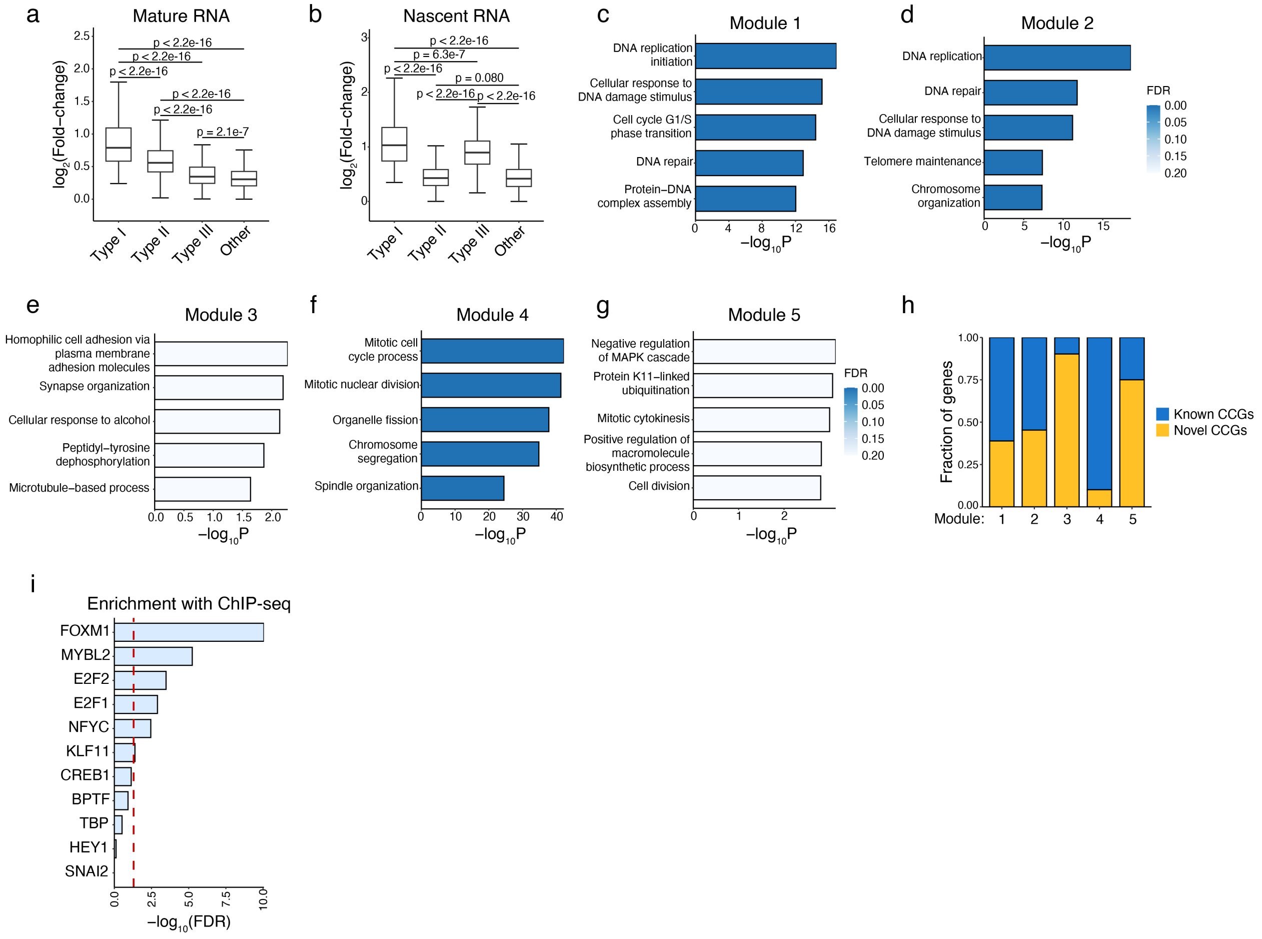


**Supplementary figure 5.** Type I CCGs in HEK293T cells. (**a**) Overall fold changes in mature RNA for different types of CCGs along cell cycle. (**b**) Overall fold changes in nascent RNA for different types of CCGs along cell cycle. (**c-g**) Gene Ontology (GO) enrichment analysis on Type I kinetic modules. (**h**) Fractions of novel CCGs in 5 kinetic modules. (**i**) Enrichment analysis between the identified targets genes and the binding targets identified by ChIP-seq for each TF. The red dashed line corresponds to -$\log_{10}(0.05)$.

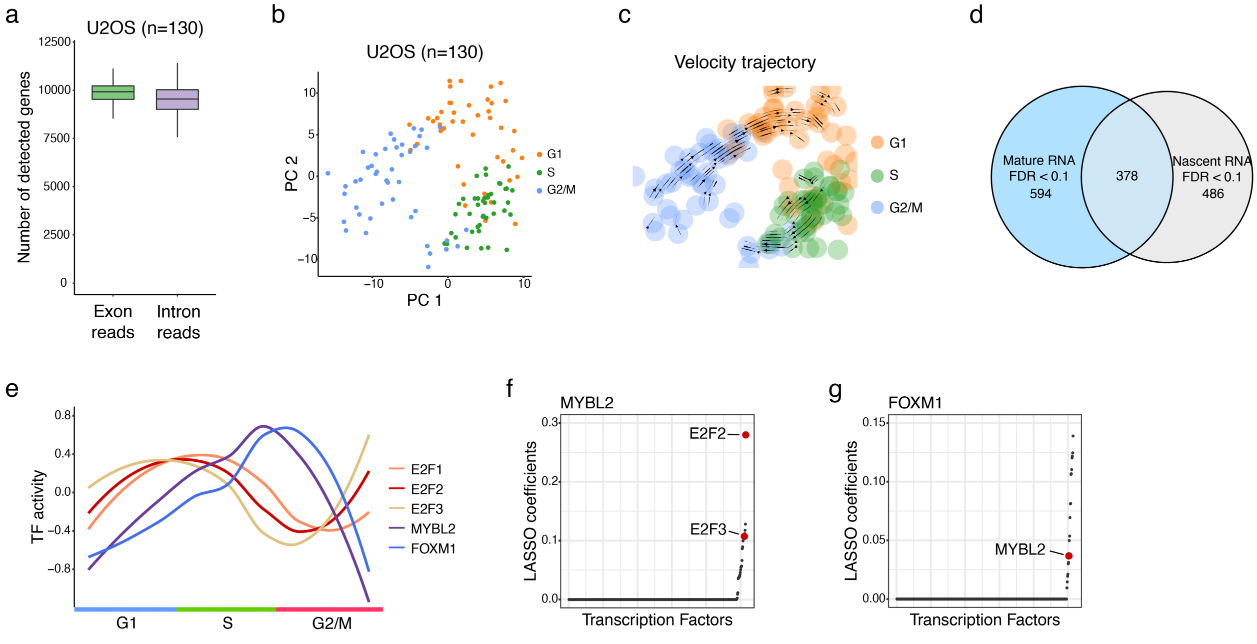


**Supplementary figure 6.** Cell cycle analysis on U2OS cell line. (**a**) The number of detected genes by exon reads or intron reads in single U2OS cells by snapTotal-seq. (**b**) PCA plot of U2OS cells. (**c**) The velocity trajectory projected by RNA velocity analysis. (**d**) Differential gene expression analyses identifying the genes with significant changes at the mature RNA or nascent RNA level along cell cycle in U2OS cell line. (**e**) The activities of different TF modules along cell cycle. (**f**) Identifying the regulatory relationships between G1 TFs and *MYBL2*. The direct regulatory links (colored in red) were identified by ChIP-seq verification. (**g**) Identifying the regulatory relationship between *MYBL2* and *FOXM1*. The direct regulatory links (colored in red) were identified by ChIP-seq verification.

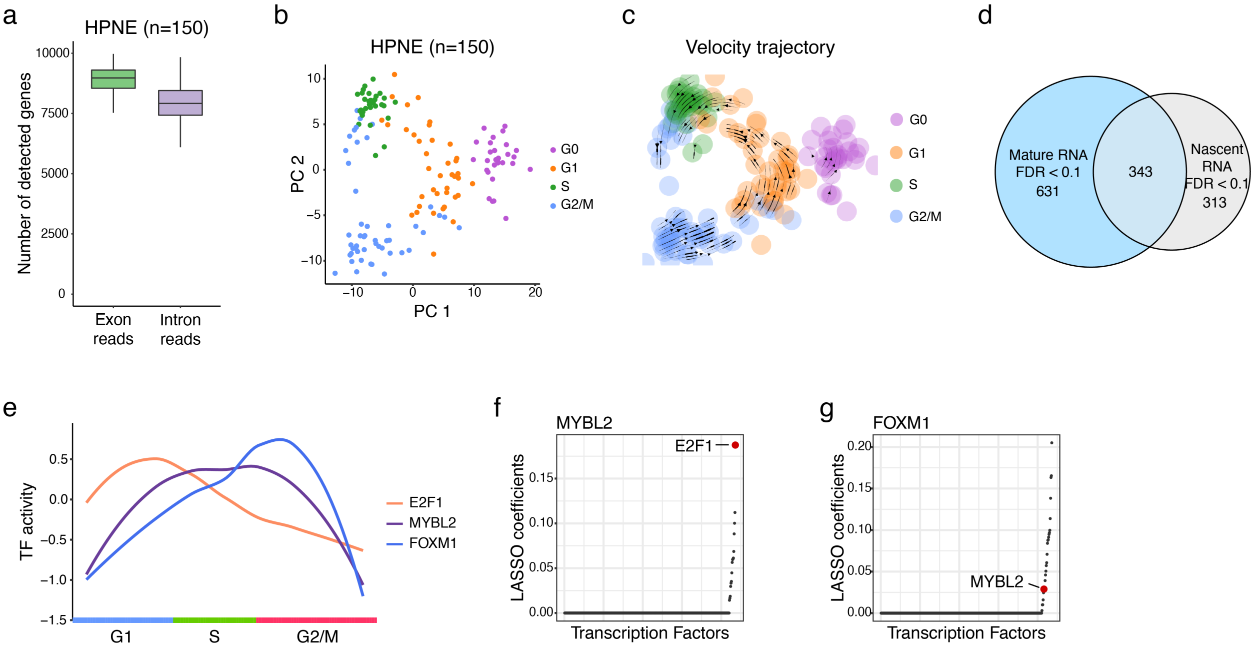


**Supplementary figure 7.** Cell cycle analysis on HPNE cell line. (**a**) The number of detected genes by exon reads or intron reads in single HPNE cells by snapTotal-seq. (**b**) PCA plot of HPNE cells. (**c**) The velocity trajectory projected by RNA velocity analysis. (**d**) Differential gene expression analyses identifying the genes with significant changes at the mature RNA or nascent RNA level along cell cycle in HPNE cell line. (**e**) The activities of different TF modules along cell cycle. (**f**) Identifying the regulatory relationships between *E2F1* and *MYBL2*. The direct regulatory links (colored in red) were identified by ChIP-seq verification. (**g**) Identifying the regulatory relationship between *MYBL2* and *FOXM1*. The direct regulatory links (colored in red) were identified by ChIP-seq verification.

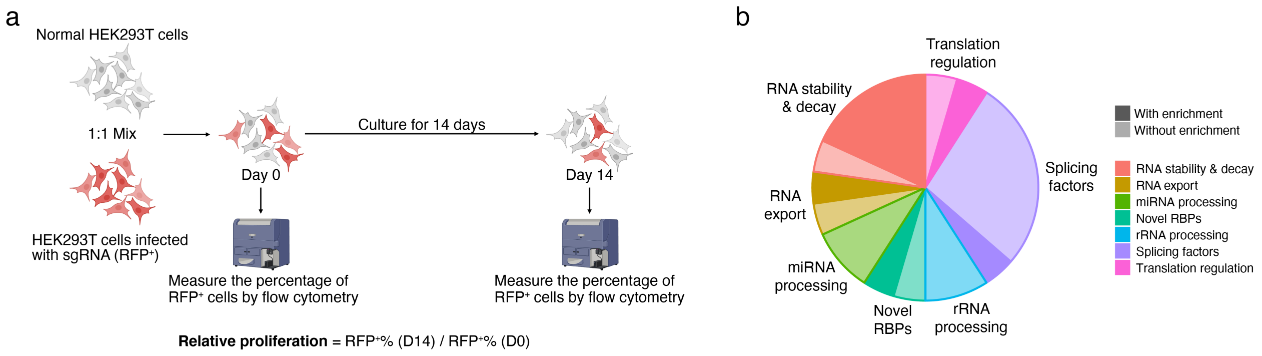


**Supplementary figure 8.** Post-transcriptional regulation in cell cycle. (**a**) The experimental scheme to examine the effects of the genes of interest on cell proliferation. (**b**) The functional categorization of RBPs with significant gene expression changes along cell cycle.

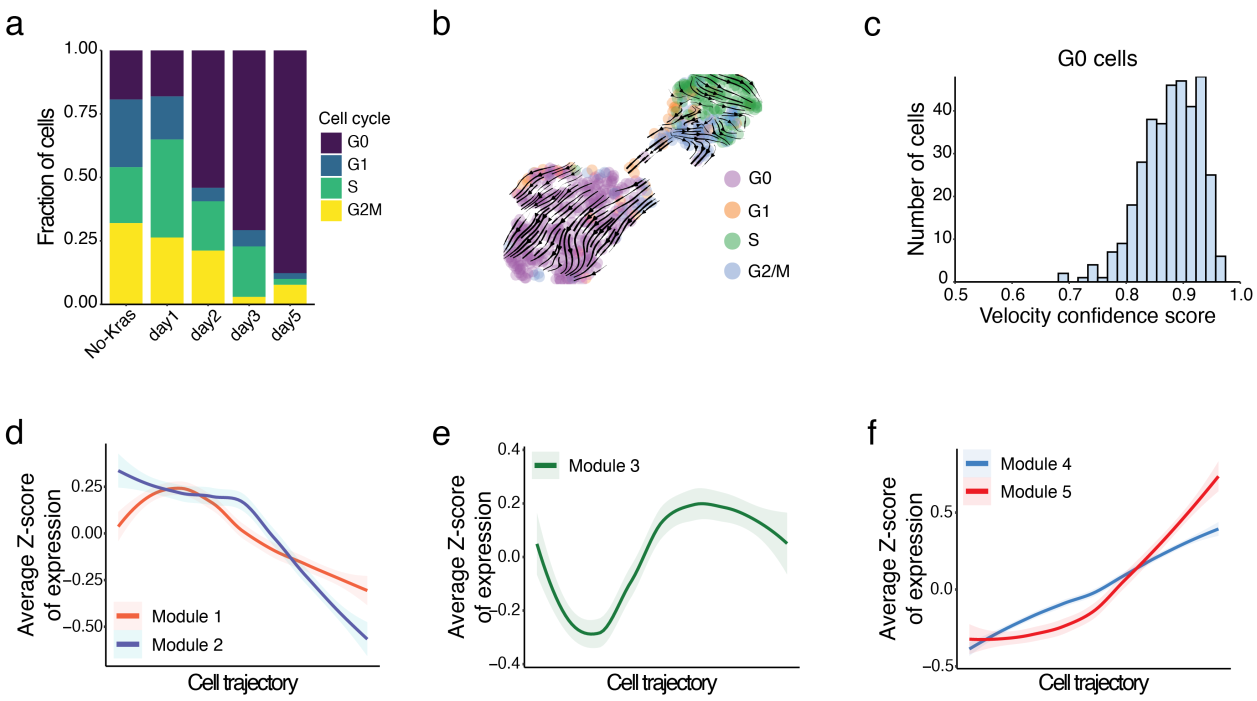


**Supplementary figure 9.** Gene expression dynamics during oncogene-induced senescence. (**a**) The fraction of cells at different cell cycle stages along the oncogene-induced senescence. (**b**) The projected velocity trajectory of oncogene-induced senescence by RNA velocity analysis. The cells are colored by the cell cycle stages. (**c**) The distribution of velocity confidence scores of all G0 cells. (**d-f**) The smoothed curves of the transcriptional dynamics along the latent time of each module. The smoothed curves were derived by using loess function. Shade, 0.95 confidence interval.

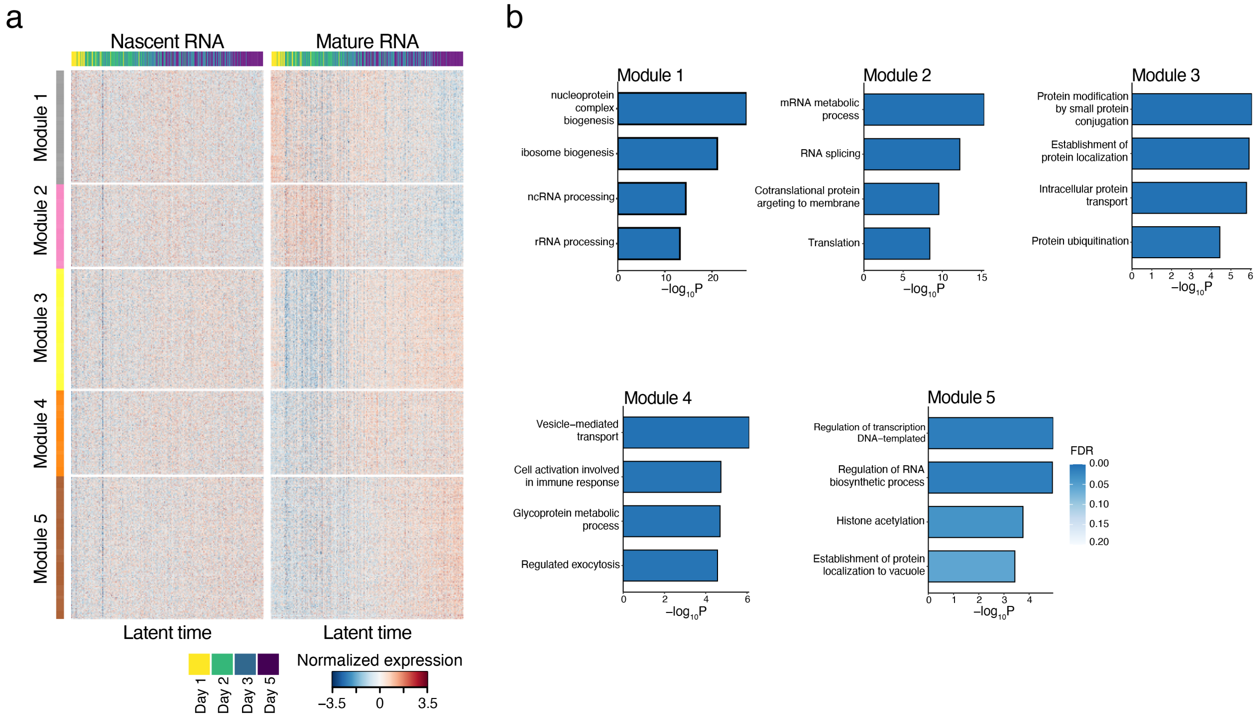


**Supplementary figure 10.** Functional analysis on Type II DEGs of oncogene-induced senescence. (**a**) The gene expression heatmap of the genes with significant changes at the mature RNA level only (Type II DEGs) along the latent time. (**b**) The GO functional enrichment for each kinetic module.

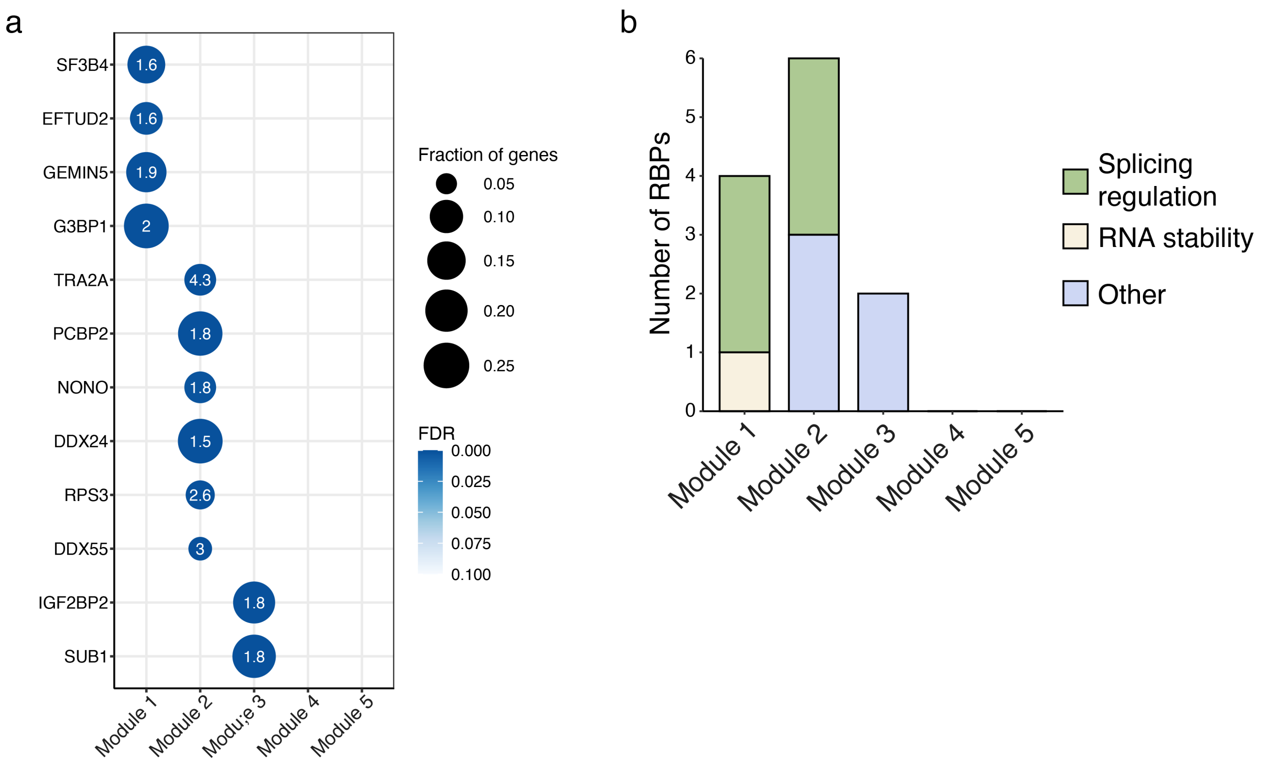


**Supplementary figure 11.** Enrichment analysis on the target genes of RBPs and Type II DEGs. (**a**) The significant enrichment of the target genes of RBPs with Type II DEGs. The RBPs need to meet the following criteria to be considered as significantly enriched: 1) The binding targets are significantly overrepresented (FDR < 0.1) in at least one of these modules; 2) The expression of the RBPs needs to be correlated (r > 0.3 or r < -0.3) with the corresponding kinetic module. (**b**) The functional categorization of RBPs with significant enrichment in Type II DEGs.

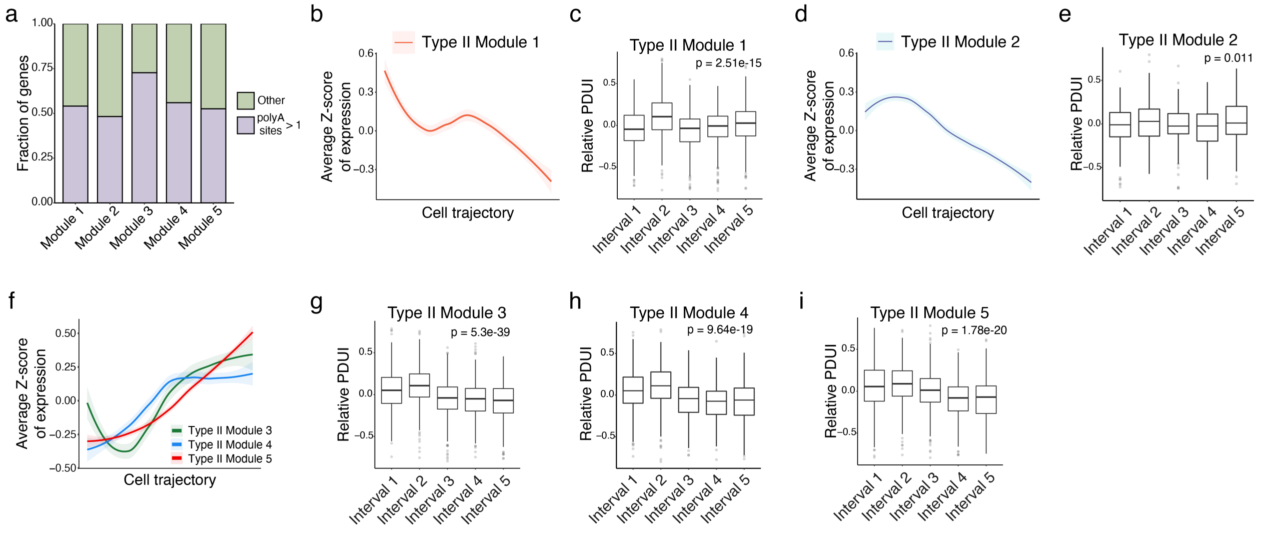


**Supplementary figure 12.** Alternative polyadenylation analysis on Type II DEGs. (**a**) The fraction of genes with multiple polyA sites in each Type II kinetic module. (**b**) The smoothed gene expression curves along the cell trajectory of oncogene-induced senescence for Type II Module 1. (**c**) The changes in polyA site usage of the genes in Type II Module 1 along the oncogene-induced senescence. (**d**) The smoothed gene expression curve along the latent time for Type II Module 2. (**e**) The changes in polyA site usage of the genes in Type II Module 2 along the oncogene-induced senescence. (**f**) The smoothed gene expression curves along the cell trajectory of oncogene-induced senescence for Type II Module 3, 4 and 5. (**g-i**) The changes in polyA site usage of the genes in Type II Module 3, 4 and 5 along the oncogene-induced senescence. PDUI: percentage of distal polyA site usage index. The relative PDUI was calculated by dividing the PDUI in each interval by the average PDUI across five intervals, followed by log-transformation. Therefore, the positive relative PDUI represents the increased usage of distal polyA sites, and the negative relative PDUI represents the decreased usage of distal polyA sites.
